# Single-cell multi-omics reveals insights into differentiation of rare cell types in mucinous colorectal cancer

**DOI:** 10.1101/2024.02.01.578409

**Authors:** Christopher A. Ladaika, Ahmed H. Ghobashi, William C. Boulton, Samuel A. Miller, Heather M. O’Hagan

**Affiliations:** Genome, Cell, and Developmental Biology, Department of Biology, Indiana University Bloomington, Bloomington, IN, 47405, USA; Medical Sciences Program, Indiana University School of Medicine, Bloomington, IN, 47405, USA; Indiana University Melvin and Bren Simon Comprehensive Cancer Center, Indianapolis, IN, 46202, USA; Department of Medical and Molecular Genetics, Indiana University School of Medicine, Indianapolis, IN, 46202, USA

## Abstract

Neuroendocrine cells have been implicated in therapeutic resistance and worse overall survival in many cancer types. Mucinous colorectal cancer (mCRC) is uniquely enriched for enteroendocrine cells (EECs), the neuroendocrine cell of the normal colon epithelium, as compared to non-mucinous CRC. Therefore, targeting EEC differentiation may have clinical value in mCRC. Here, single cell multi-omics was used to uncover epigenetic alterations that accompany EEC differentiation, identify STAT3 as a novel regulator of EEC specification, and discover a rare cancer-specific cell type with enteric neuron-like characteristics. Further experiments demonstrated that lysine-specific demethylase 1 (LSD1) and CoREST2 mediate STAT3 demethylation and regulate STAT3 chromatin binding. Knockdown of CoREST2 in an orthotopic xenograft mouse model resulted in decreased primary tumor growth and lung metastases. In culmination, these results provide rationale for new LSD1 inhibitors that target the interaction between LSD1 with STAT3 or CoREST2, which may improve clinical outcomes for patients with mCRC.

---

Across different cancer types, intratumor heterogeneity has been shown to promote tumor progression, metastasis, and resistance to therapy^1, 2^. Intratumor heterogeneity is influenced by many factors, including genetic variability, epigenetic alterations, and changes in the tumor microenvironment^1, 2^. One manifestation of heterogeneity in cancer is the presence of distinct cell types that can be advantageous to cancer growth and survival^2^. For instance, in ovarian cancer and glioblastoma, cancer stem cells are key drivers of therapeutic resistance and cancer recurrence^3, 4^.

Neuroendocrine cells are specialized cells that exhibit characteristics of neurons and endocrine cells^5^. These cells are found in normal tissue and recent work has highlighted the importance of neuroendocrine cells in several cancer types^6–9^. Neuroendocrine cells can form cancers called neuroendocrine carcinomas or exist within a heterogeneous cancer population^5^. Neuroendocrine cells have been shown to play a role in tumor aggressiveness and therapeutic resistance in breast, lung, and prostate cancer^10–12^.

Colorectal cancer is the second leading cause of cancer-related deaths in men and women combined^13^. Accounting for approximately 20% of colorectal cancer cases, mucinous colorectal cancer (mCRC) is characterized by tumors with mucus accounting for at least 50% of the tumor volume^13, 14^. Compared to non-mCRC, mCRC has been reported to be more common in women, located in the proximal colon, and have worse overall survival^13, 14^. Previously, our lab has shown that enteroendocrine cells (EECs), the neuroendocrine cells of the large intestine, are enriched in mCRC and promote cancer cell survival via the secretion of pro-survival factors^6^. Additionally, previous studies have found an association between levels of EEC-secreted factors and worse overall survival^15^. These findings indicate that identifying factors that drive EEC specification in mCRC will have clinical relevance.

In this study, we use single cell multi-omics to uncover new insights into epigenetic changes and to identify novel potential regulators of EEC specification in mCRC. We then experimentally show that signal transducer and activator of transcription 3 (STAT3) acts with lysine specific demethylase 1 (LSD1) to promote EEC differentiation in mCRC. To carry out its enzymatic function, LSD1 typically functions within a larger transcriptional regulatory complex, such as the CoREST complex^16–19^. In this study, we demonstrate that CoREST2, one of the three CoREST protein paralogs, is primarily expressed within the EEC lineage, is critical for EEC specification, interacts with and regulates STAT3, and decreases tumor progression and lung metastasis in an orthotopic xenograft mouse model. Collectively, these findings improve our understanding of EEC differentiation in mCRC and suggest CoREST2 as a novel therapeutic target.

## Results

### Single Cell Multi-omics Reveals Chromatin Changes During EEC Differentiation

To study EEC differentiation in mCRC, we utilized the small molecule drug ISX9, which promotes the differentiation of EECs in the normal colon^20^. Using the mCRC cell line HT29, we verified that ISX9 promotes EEC differentiation in mCRC via immunofluorescence of EEC-promoting transcription factor INSM1 (Fig 1A). While many transcription factors that regulate EEC lineage specification have been discovered, little is known about the chromatin changes that occur during this process. To investigate the epigenetic alterations and transcription factors involved in EEC differentiation, we treated cells with ISX9 and conducted a single cell multi-omics experiment that combines single cell RNA sequencing with single cell ATAC sequencing.

**Figure 1.**
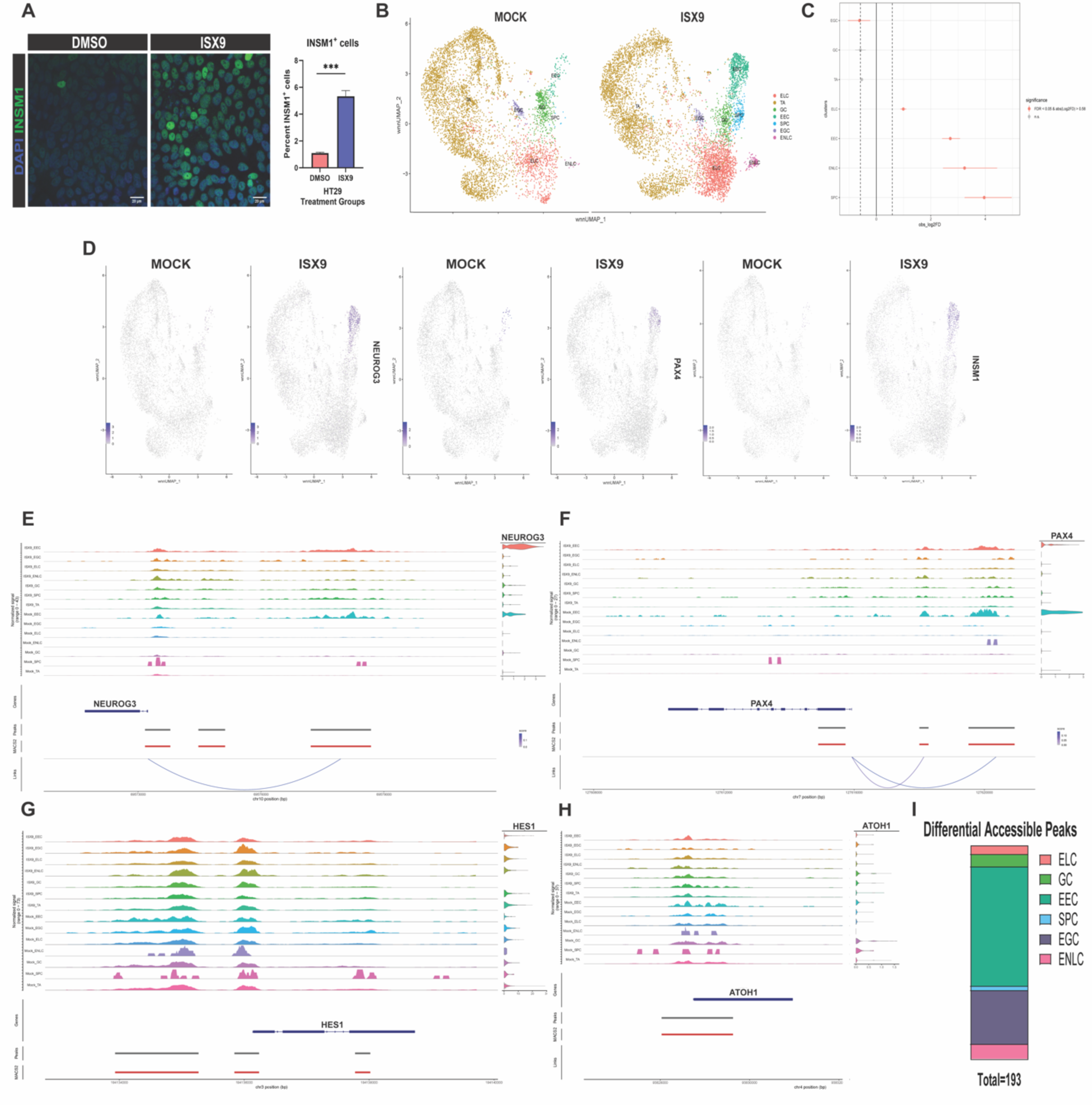
Single cell multi-omics reveals chromatin changes during EEC differentiation. **A**. Representative immunofluorescence images of EEC marker INSM1 (green) in HT29 cells treated twice for 24hrs with DMSO or ISX9 (40 µM). Graph shows quantification of percent INSM1 positive cells +/- SEM (N=3). **B.** UMAP dot plot of HT29 single cell multi-omics samples treated with DMSO (MOCK, left) or ISX9 (right) treated as in A. Samples are colored by cell type/cluster. TAs are transit amplifying cells, ELCs are enterocyte-like cells, SPCs are secretory progenitor cells, EGCs are early goblet cells, GCs are goblet cells, EECs are enteroendocrine cells, and ENLCs are enteric neuron-like cells **C.** Relative cell proportions for each cluster in DMSO vs ISX9 treated HT29 cells. Red dots have FDR < 0.05 and mean absolute |log_2_FD| > 0.58 in ISX9 vs. MOCK treated cells. **D.** Feature blots of normalized expression values of EEC marker genes *NEUROG3*, *PAX4*, and *INSM1.* **E.** Peaks to genes linkage map of NEUROG3. Peaks show ATAC-seq peak accessibility at *NEUROG3* regulatory regions, violin plot shows expression of *NEUROG3*, curved blue line shows significant correlation between peak accessibility and expression of *NEUROG3*. **F-H.** Peaks to genes linkage map of *PAX4, HES1,* and *ATOH1*. Visualization is the same as in E. **I.** Number of differential accessible peaks in Mock and ISX9-integrated specific cell type cluster compared to all other clusters. All clusters are included except the TA cluster.

Cells were clustered based on transcriptomic and chromatin accessibility data and clusters were annotated through marker analysis (Fig 1B and Extended Data Fig 1). As expected, treatment with ISX9 increased the proportion of EECs, which were defined by high expression of known EEC-promoting transcription factors (Fig 1C-1D). To determine chromatin changes that accompany EEC differentiation, we performed a peak-to-gene linkage analysis of genes encoding EEC-promoting transcription factors. At *NEUROG3*, accessibility at the regulatory region 5000 base pairs upstream of the transcriptional start site showed the most correlation with expression of *NEUROG3* (Fig 1E). Meanwhile, both the regulatory regions located 2000 and 4000 base pairs upstream of the *PAX4* transcriptional start site had a significant correlation between chromatin accessibility and *PAX4* expression (Fig 1F). Interestingly, ISX9 treatment increased chromatin accessibility at regulatory regions of EEC-promoting transcription factors in non-EEC cell types without inducing expression of these genes (Fig 1E-1F and Extended Data Fig 2A-2B). For example, in Mock-treated cells, the chromatin is only accessible at denoted regulatory regions of *NEUROG3* and *PAX4* in the secretory progenitor and EEC clusters; however, with ISX9 treatment, the chromatin at these regions becomes accessible in all clusters (Fig 1E-1F).

Dynamic changes in chromatin accessibility were most prominent in the EEC lineage, as we did not observe changes in chromatin accessibility of non-EEC lineage-promoting genes in different clusters. For example, *HES1* and *ATOH1*, which regulate the bifurcation of the absorptive and secretory lineages in the colon^21^, had relatively equal chromatin accessibility in the different clusters, regardless of treatment (Fig 1G and 1H). Similar trends were also observed for secretory progenitor markers *DLL1* and *DLL4*, early goblet cell marker *REG4*, and goblet cell markers *SPDEF* and *MUC2* (Extended Data Fig. 2C-2G). This finding is further corroborated by determining that EECs had the most differentially accessible peaks of all clusters other than the transit amplifying (TA) cluster (Fig 1I and Extended Data Fig 3). Collectively, these results show that dynamic changes in chromatin accessibility at genes encoding EEC-promoting transcription factors are important for EEC differentiation.

### Identification of Rare Enteric Neuron-Like Tumor Cells

During cluster annotation, one cluster did not fit the description of any cell type found in the normal colon epithelium. This cluster expressed several strong marker genes, including *AKAP12*, *THBS2*, and *PRSS23*, but did not express markers of secretory cells (Fig 2A-2B). To annotate this cluster, we submitted the top 100 genes based on adjusted p-value with increased expression in this cluster relative to all other clusters to the database *Enrichr*^22^. Intriguingly, five of the top 10 gene ontology biological processes were related to neuron-related processes (Fig 2C). Submission of this gene list to the Descartes Cell Atlas returned results of various neuronal cells, including enteric neurons of the pancreas and stomach (Fig 2C). Given these results, we annotated this cluster as enteric neuron-like cells (ENLCs) (Fig 1B). Using AKAP12 immunofluorescence in our mCRC cell line the percentage of AKAP12-positive ENLCs was approximately 0.2% at baseline and increased to 1.0% after ISX9 treatment, matching the ENLC percentages in our multi-omics data (Fig 2D). Next, we performed immunohistochemistry staining for AKAP12 on a tissue microarray containing colon adenocarcinoma and adjacent normal tissue patient samples. Interestingly, while in 37% of samples the percentage of AKAP12-positive tumor cells was between 1-10%, reflecting AKAP12 being a marker for a rare cell type; in 12% of samples 25-90% of tumor cells were AKAP12-positive (Fig 2E). Strikingly, there were no AKAP12-positive normal colon epithelial cells in any of the samples examined, suggesting the ENLCs emerge during carcinogenesis (Fig 2E).

**Figure 2.**
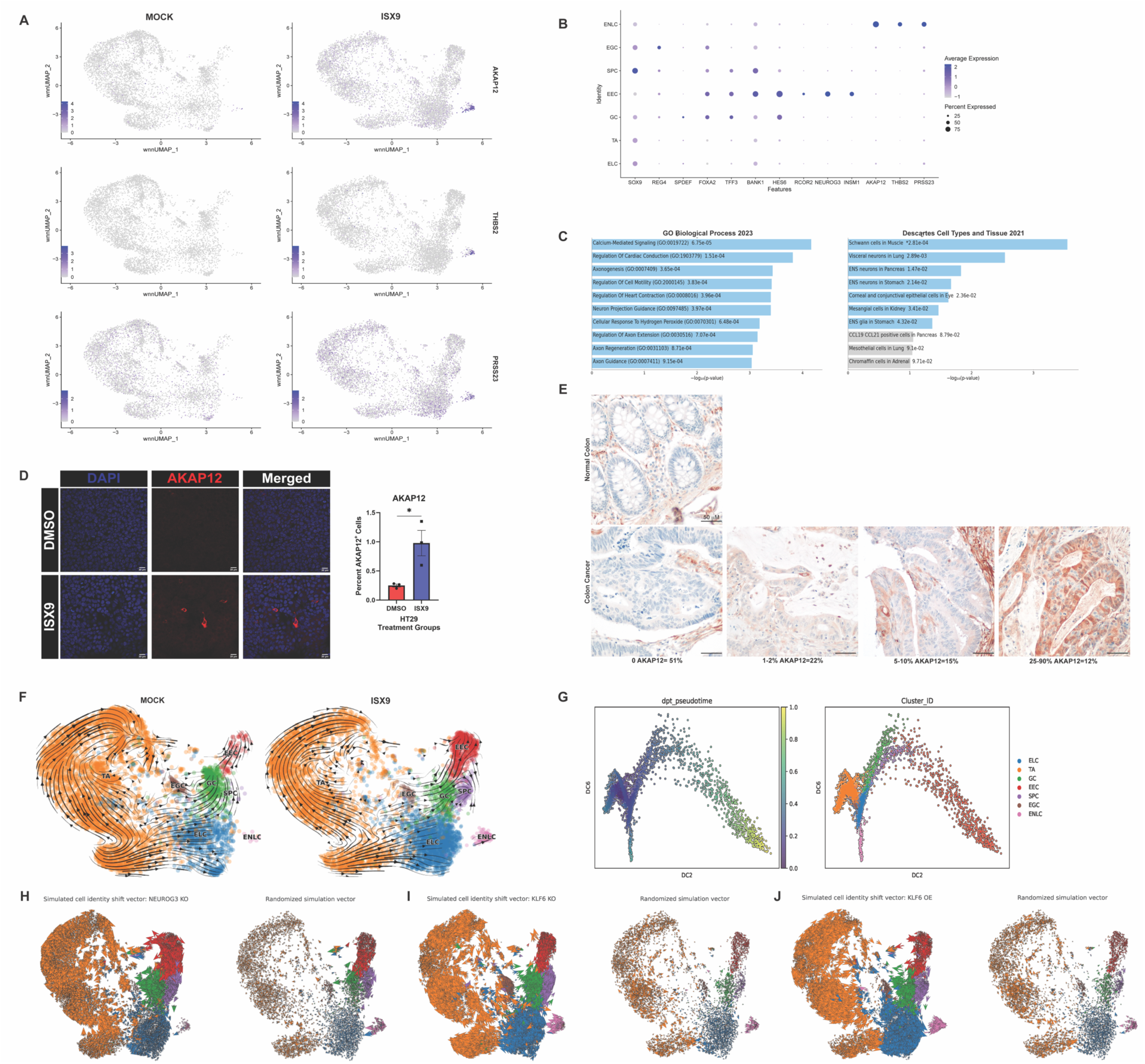
Enteric neuron-like cells are a rare non-secretory cell type in mucinous colorectal cancer. **A.** UMAP dot plots of normalized expression values of enteric neuron-like cell (ENLC) marker genes *AKAP12*, *THBS2*, and *PRSS23* in HT29 cell scRNA-seq data from Figure 1. **B.** Dot plot showing various marker genes expression in ISX9 treated samples across all annotated cell types. The size of the dot is proportional to the percentage of cells that express a given gene and the color scale indicates the average scaled gene expression within the specific cell population. **C.** Gene Ontology (Left) and Descartes Cell Type and Tissue (Right) analyses of top 100 genes differential expressed in ENLC cluster relative to all other clusters based on adjusted p-value. **D.** Representative immunofluorescence images of ENLC marker AKAP12 (red) in HT29 cells treated with two 24hr treatments of DMSO or ISX9 (40 µM). Graph shows quantification of percent AKAP12-positive cells +/- SEM (N=3). **E.** Representative AKAP12 immunohistochemistry staining of normal colon (top) and colon tumors (bottom) from human colon tumor microarray. **F.** RNA velocity stream plots of MOCK and ISX9 treated cells. **G.** (Left) Diffusion map of ISX9 treated cells arranged in diffusion pseudotime; (Right) annotated diffusion map. **H.** RNA velocity plot showing cell identity shift following *NEUROG3* simulated knockout (left) and randomized simulated vector (right). **I.** RNA velocity plot showing cell identity shift following *KLF6* simulated knockout (left) and randomized simulated vector (right). **J.** RNA velocity plot showing cell identity shift following *KLF6* simulated overexpression (left) and randomized simulated vector (right). Significance was determined by Student’s t-test **(D).** *P≤ 0.05.

### RNA Trajectory and Diffusion Mapping Analyses Provide Insights into ENLC Differentiation

To ascertain how ENLCs are related to other cell types, we performed an RNA velocity trajectory analysis (Fig 2F). In the Mock-treated group, EECs primarily emerge from secretory progenitors; however, in the ISX9-treated group, the trajectories of the TA, early goblet cell, and goblet cell clusters reoriented toward the EEC cluster (Fig 2F). Notably, the ENLCs did not originate from the enterocyte-like cluster or secretory progenitor cluster, suggesting ENLCs do not belong to the absorptive or secretory lineages (Fig 2F). To further evaluate the differentiation of ENLCs, we conducted a diffusion mapping analysis, whereby cells are arranged by the probability of differentiating into different cell types. Consistent with our trajectory analysis, we were able to capture that ENLCs follow a different trajectory from the secretory lineage by using diffusion components 2 and 6 (Fig 2G). Evaluation of cell marker expression patterns across differentiation revealed that ENLC markers were upregulated only during the differentiation of ENLCs, while secretory progenitor and EEC markers were not expressed in the ENLC cluster (Fig 2B and Extended Data Fig 5A-5B). Collectively, this data shows ENLCs are a rare non-secretory cell type.

### Gene Regulatory Network Analysis Identifies Candidate Regulators of ENLC Specification

To further explore the regulation of ENLCs, we used the ISX9-treated samples from our multi-omics data to construct cell type specific gene regulatory networks (GRNs) (Extended Data Fig 4). Candidate regulators were identified through a centrality analysis, which estimates the importance of a gene in the regulation of a particular cell fate. Based on betweenness centrality, KLF6 ranked as the top regulator of the ENLC fate (Extended Data Fig 5C). To evaluate how KLF6 impacts ENLC differentiation, we used CellOracle to perform *in silico* perturbations of ISX9-treated cells. As proof of principle, we simulated the knockout of *NEUROG3*, which is essential for EEC differentiation^23^, and observed a transition of cells away from the EEC fate (Fig 2H). Interestingly, simulated knockout of KLF6 transitioned cells away from the ENLC cluster and towards the EEC cluster (Fig 2I). Conversely, simulated overexpression of KLF6 increased ENLC specification, and decreased EEC differentiation (Fig 2J). Together, these results suggest that KLF6 promotes ENLC differentiation and represses EEC differentiation. Based on our centrality analysis, another top potential regulator of ENLCs was FOSL1. FOSL1 stood out, as a motif analysis of differential accessible peaks in the ENLC cluster identified several AP-1 transcription factors associated motifs, including FOSL1 and JUN (Extended Data Fig 5D). Simulated knockout of FOSL1, indicates FOSL1 is predicted to repress both the ENLC and EEC fate (Extended Fig 5E). Collectively these results reveal new potential regulators of two rare cell types in mCRC.

### STAT3, JAK2, and Calcium Promote EEC Specification

Based on our multi-omics data, while the opening of chromatin at EEC-promoting transcription factors is important for promoting differentiation, it is insufficient for driving expression (Fig 1E-1F). Therefore, we set out to identify another transcription factor that was driving expression of these genes. To computationally determine transcription factors associated with EEC differentiation, we performed a differential motif accessibility analysis comparing EECs to all other clusters. Several transcription factors known to be associated with EEC and/or neuroendocrine cell differentiation were identified, including INSM1 and SOX4 (Extended Data Fig 3)^24–25^. Another motif enriched in the EEC cluster of accessible peaks was STAT3 (Figure 3A and Extended Data Figure 3). STAT3 stood out to us as STAT3 promotes the differentiation of pancreatic beta cells and mouse spermatogonia cells through promoting *NEUROG3* expression^26–28^. Additionally, STAT3 has been shown to promote the neuroendocrine cell fate in androgen receptor-independent prostate cancer^29^. Based on the relevance of STAT3 in these contexts, we decided to further evaluate the role of STAT3 in regulating EEC differentiation in mCRC.

**Figure 3.**
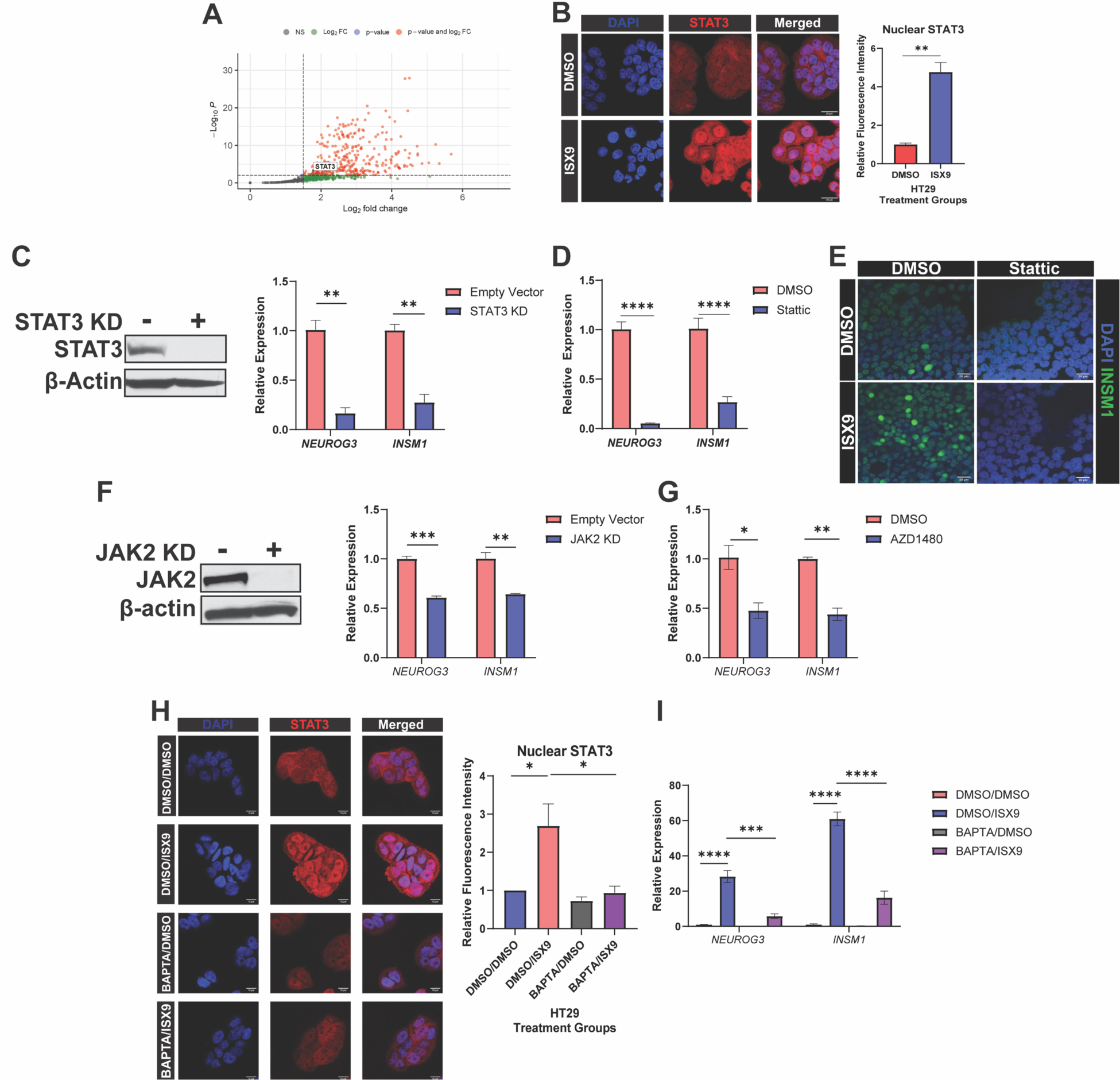
STAT3 promotes EEC differentiation in mucinous colorectal cancer. **A.** Volcano plot showing transcription factor motif enrichment in differential accessible peaks in EECs vs non-EEC clusters. Red dots have a log2FC > 1.5 and p-value < 0.05. **B.** Representative immunofluorescence showing increased nuclear STAT3 (red) in HT29 cells treated with vehicle or ISX9 (40 µM) for 4hrs. Graph shows quantification of nuclear STAT3 +/- SEM (N=3). **C.** (Left) Western blot showing shRNA mediated knockdown of STAT3. (Right) qRT-PCR of EEC marker RNA expression in empty vector and STAT3 knockdown (STAT3 KD) HT29 cells. **D.** qRT-PCR of EEC markers after treating HT29 cells twice for 24hrs with vehicle or STAT3 inhibitor (Stattic, 10 µM). **E.** Representative immunofluorescence images of EEC marker INSM1 (green) in HT29 cells treated for 24hrs with vehicle or Stattic (10 µM) prior to two 24hr treatments with vehicle or Stattic (10 µM) and vehicle or ISX9 (40 µM). **F.** (Left) Western blot showing shRNA mediated knockdown of JAK2 in HT29 cells. (Right) qRT-PCR of EEC markers in empty vector and JAK2 knockdown (JAK2 KD) HT29 cells. **G.** qRT-PCR of EEC markers after treating HT29 cells twice for 24hrs with vehicle or JAK2 inhibitor AZD-1480 (5 µM). **H.** qRT-PCR of EEC markers following pre-treatment of HT29 cells with vehicle or calcium chelator BAPTA-AM (20 µM) twice for 1hr prior to treatment with vehicle or ISX9 (40 µM) for 24hrs. **I.** Representative immunofluorescence images of STAT3 (red) in HT29 cells treated with vehicle or BAPTA-AM (20 µM) for 1hr prior to treatment with vehicle or ISX9 (40 µM) for 4hrs. Graph shows quantification of nuclear STAT3 +/- SEM (N=3). Significance was determined by Student’s t-test **(B,C, D, F, and G)** and one-way ANOVA with Tukey pairwise multiple comparison testing (**H and I).** *P≤ 0.05, ** P ≤ 0.01, *** P ≤ 0.001, **** P ≤ 0.0001.

ISX9 treatment increased STAT3 nuclear localization in HT29 cells and increased levels of phosphorylated STAT3^Y705^ in normal colon organoids. (Fig 4B and Extended Data Fig 6A), suggesting that STAT3 is activated during EEC differentiation. Knocking down STAT3 or STAT3 inhibition in HT29 cells decreased the baseline expression of EEC markers *NEUROG3* and *INSM1* (Fig 3C-3D and Extended Data Fig 6B-6C). These results were corroborated in SW403 and LS174T cell lines and in normal colon organoids (Extended Data Fig 6D-6F). STAT3 inhibition also decreased expression of secretory marker *ATOH1* but did not affect expression of absorptive, stem, or goblet cell markers, suggesting that STAT3 specifically promotes EEC differentiation (Extended Data Fig 6G). Additionally, immunofluorescence for INSM1 indicated that STAT3 inhibition dramatically decreased the percentage of EECs both at baseline and in combination with ISX9 (Fig 3E). We also found that knocking down or inhibiting STAT3-activating kinase JAK2 decreased expression of EEC markers in HT29 and NCI-H508 cells (Fig 3F-3G and Extended Data Fig 6H-6I). STAT3 can be activated by various signals including calcium. Our multi-omics experiment revealed many genes associated with calcium-dependent signaling were highly expressed in EECs (Extended Data Fig 6J-6K). Pre-treatment with calcium chelator *BAPTA-AM* prior to ISX9 treatment, blocked the ISX9-induced increase in nuclear STAT3 and attenuated ISX9-meditated EEC differentiation (Fig 3H-3I). Collectively, these results show that calcium and JAK2 activate STAT3 to promote EEC differentiation.

**Figure 4.**
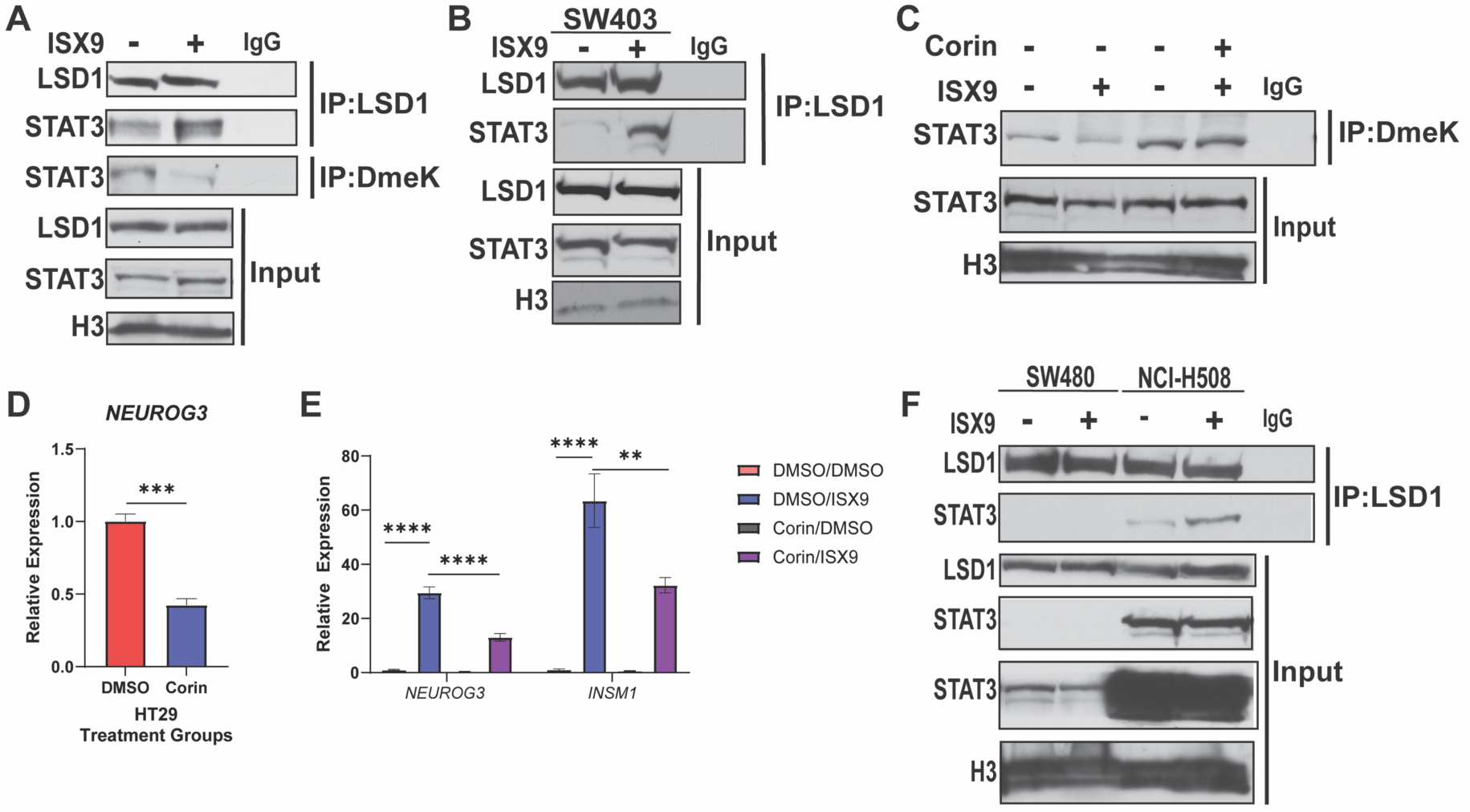
LSD1 interacts with and demethylates STAT3 to promote EEC differentiation. **A.** Western blot of LSD1 immunoprecipitation (IP) and Dimethyl-lysine (DmeK) IP performed using nuclear lysates prepared from HT29 cells treated with vehicle or ISX9 (40 µM) for 4hrs. IgG IP serves as a negative control. **B.** Western blot of LSD1 IP performed using nuclear lysates prepared from SW403 cells treated as in A. IgG IP serves as a negative control. **C.** Western blot of DmeK IP performed using nuclear lysates prepared from HT29 cells treated with vehicle or Corin (50 nM) for 24hrs prior to treatment with DMSO or Corin (50 nM) and DMSO or ISX9 (40 µM) for 4hrs. IgG IP serves as a negative control. **D.** qRT-PCR of EEC marker RNA expression after treating cells with vehicle or LSD1 inhibitor (Corin, 50 nM) for 24hrs. **E.** qRT-PCR of EEC markers after treating cells with vehicle or Corin (50 nM) for 24hrs prior to two 24hr treatments of vehicle or Corin (50 nM) and DMSO or ISX9 (40 µM). **F.** Western blot of LSD1 IP performed using nuclear lysates prepared from SW480 (non-mucinous) or NCI-H508 (mucinous) cells treated as in A. Significance was determined by Student’s t-test **(D)**, one-way ANOVA with Tukey pairwise multiple comparison testing **(E**). ** P ≤ 0.01, *** P ≤ 0.001, **** P ≤ 0.0001.

### LSD1 Demethylates STAT3 to Promote EEC Differentiation

STAT3 can be regulated by various post-translational modifications such as phosphorylation and methylation^29^. Previous studies have separately linked LSD1 to promoting EEC differentiation and demethylating STAT3^6, 30^. To test whether LSD1 demethylates STAT3 during EEC differentiation, we treated cells with ISX9 and performed LSD1 and dimethyl-lysine immunoprecipitations. ISX9 increased the interaction between LSD1 and STAT3 and decreased the levels of dimethyl-lysine STAT3 in multiple cell lines (Fig 4A-4B and Extended Data Fig 7A-7B). The demethylation of STAT3 was blocked using dual LSD1 and HDAC1/2 inhibitor Corin, demonstrating that LSD1 is demethylating STAT3 (Fig 4C). Corin treatment also decreased expression of EEC marker genes at baseline and following ISX9 treatment (Fig 4D-4E). Fitting with a specific role in promoting EEC differentiation, no interaction between LSD1 and STAT3 was detected in the non-mucinous cell line SW480, which is regarded as primarily stem-like and lacks EECs (Fig 4F). Collectively, these data show that LSD1 likely promotes EEC differentiation by demethylating STAT3.

### CoREST2 is Expressed Primarily in EECs and Promotes EEC Differentiation

Three CoREST paralogs can interact with LSD1 in the CoREST complex^31^. Previously, our lab performed single cell RNA-seq on HT29 and NCI-H508 cells and integrated the data with published single cell RNA-seq data from the normal colon^6^. Interestingly, both in that data set and our multi-omics, *RCOR1* and *RCOR3* (CoREST1 and CoREST3) were ubiquitously expressed, while *RCOR2* (CoREST2) was expressed primarily within the EEC lineage (Fig 5A and Extended Data Fig 8A-8C). Additionally, *RCOR2* regulatory regions were most highly accessible in the EEC cluster in Mock-treated cells (Extended Data Fig 8D). Immunofluorescence for EEC marker βIII-TUBULIN (β3T)^32, 33^ following individual knockdowns of CoREST paralogs, revealed that CoREST1 and CoREST3 do not affect EEC differentiation, while knocking down CoREST2 dramatically decreased the number of EECs and expression of EEC marker genes (Fig 5B-C and Extended Data Fig 8E-8F). Of note, 91.23% of CoREST2-positive cells were also β3T-positive, suggesting that most CoREST2-positive cells are EECs (Fig 5B). Like LSD1, there is an increased interaction between CoREST2 and STAT3 after ISX9 treatment and CoREST2 KD blocked the ISX9-induced decrease in levels of dimethyl-lysine STAT3 (Fig 5D-E). Notably, CoREST1 KD did not affect STAT3 dimethyl-lysine levels after ISX9 treatment (Extended Data Fig 8G). Surprisingly, overexpressing CoREST2 -FLAG did not increase expression of EEC markers (Fig 5F). However, LSD1 KD decreased CoREST2-FLAG protein levels (Extended Data Fig 8H), leading us to hypothesize that there was insufficient LSD1 to stabilize the overexpressed CoREST2-FLAG. To test this, CoREST1 was knocked down in CoREST2-FLAG overexpressing cells and a CoREST2-FLAG immunoprecipitation was performed following ISX9 treatment. CoREST1 KD increased the interaction of CoREST2-FLAG with LSD1 and increased CoREST2-FLAG protein levels (Fig 5G). There was also a slight increase in the interaction between CoREST2 and STAT3 in CoREST1 KD cells (Fig 5G). Overexpressing CoREST2 and knocking down CoREST1 had a synergistic effect on the expression of EEC markers (Fig 5H). Collectively, these results show that CoREST2 is important in EEC specification, likely by regulating STAT3 demethylation, through its interaction with LSD1.

**Figure 5.**
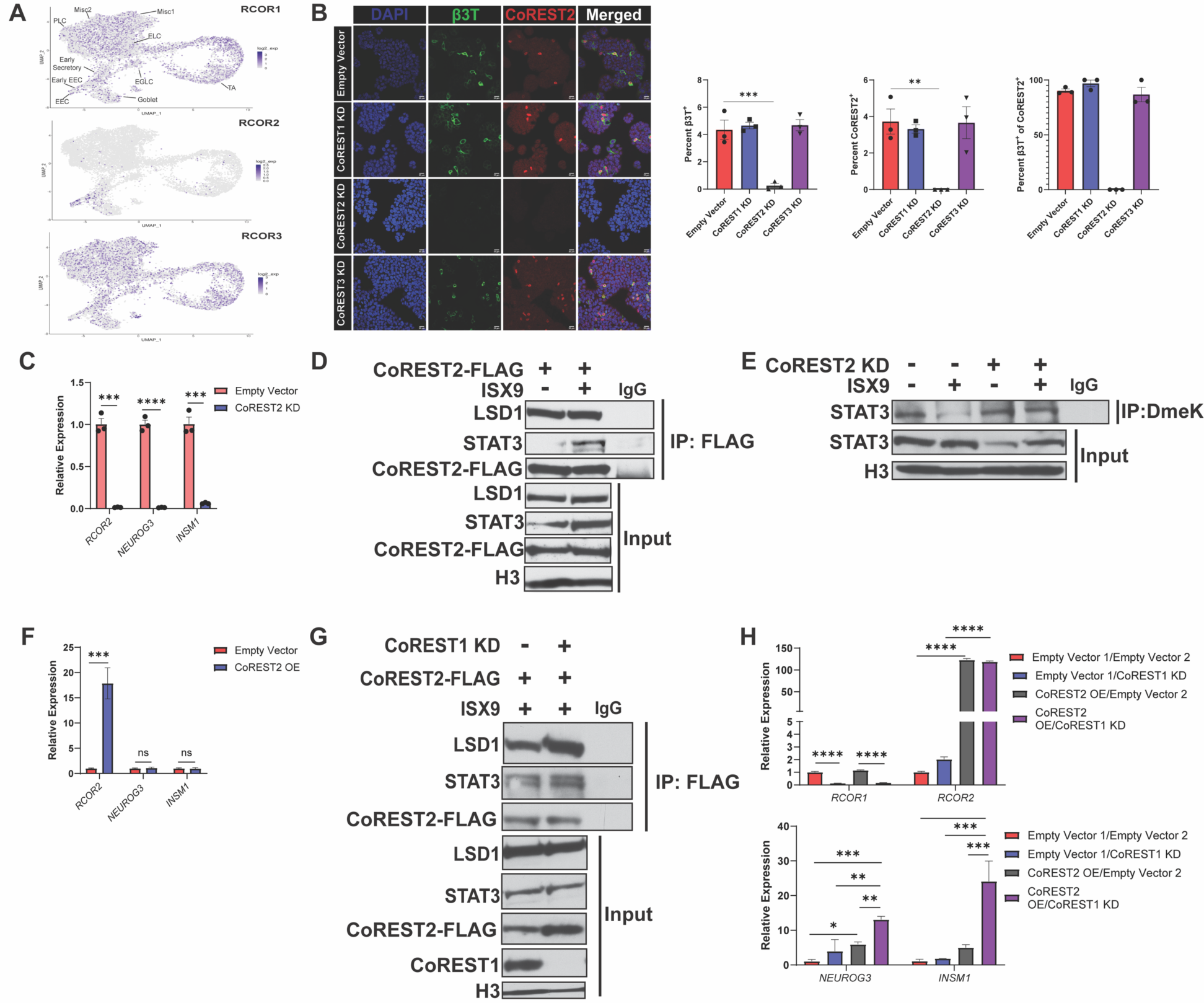
CoREST2 interacts with LSD1 and STAT3 to promote EEC differentiation. **A.** Individual UMAP plots showing the expression of *RCOR1*, *RCOR2*, or *RCOR3* in individual cells from normal colon, HT29, and NCI-H508 scRNA-seq samples. **B.** Representative immunofluorescence images of EEC marker BETA-3-TUBULIN (β3T, green) and CoREST2 (red) in empty vector, CoREST1 knockdown (KD), CoREST2 KD, and CoREST3 KD HT29 cells. Graphs show quantification of percent B3T-positive (left), percent CoREST2-positive (middle), and percent β3T-positive of CoREST2-positive cells (right). **C.** qRT-PCR of *RCOR2* and EEC marker RNA expression in empty vector and CoREST2 KD cells. **D.** Western blot of FLAG immunoprecipitation (IP) performed using nuclear lysates prepared form HT29 cells transduced with a plasmid containing FLAG tagged CoREST2 overexpression (CoREST2 OE FLAG) and treated with vehicle or ISX9 (40 µM) for 4hrs. IgG IP serves as a negative control. **E.** Dimethyl-lysine (DmeK) IP performed using nuclear lysates prepared from empty vector and CoREST2 KD HT29 cells treated with vehicle or ISX9 (40 µM) for 4hrs. **F.** qRT-PCR of *RCOR2* and EEC markers in HT29 cells transduced with an empty vector or CoREST2 OE FLAG plasmid. **G.** Western blot of FLAG IP performed using nuclear lysates prepared form empty vector and CoREST1 KD HT29 cells transduced with CoREST2 OE FLAG plasmid and treated with ISX9 (40 µM) for 4hrs. IgG IP serves as a negative control. **H.** qRT-PCR of *RCOR1* and *RCOR2* (top) and EEC markers (bottom) in empty vector and CoREST1 KD HT29 cells transduced with an empty vector or CoREST2 OE FLAG plasmid. Significance was determined one-way ANOVA with Tukey pairwise multiple comparison testing **(B and G)** and by Student’s t-test **(C).** *P≤ 0.05, ** P ≤ 0.01, *** P ≤ 0.001, **** P ≤ 0.0001.

### LSD1 and CoREST2 Regulate STAT3 Chromatin Binding

To determine where STAT3 is binding to promote EEC differentiation, STAT3 CUT&RUN was performed. ISX9 treatment increased enrichment of STAT3 at multiple sites across the genome (Fig 6A). Gene ontology analysis on STAT3 peaks from ISX9-treated cells revealed an enrichment of peaks at genes associated with neurons (Fig 6B and Extended Data Fig 9). These results support a role for STAT3 in promoting EEC differentiation, as many genes involved in neuronal differentiation also have roles in EEC differentiation. Next, to determine if CoREST2 binds to similar genes as STAT3, we performed FLAG CUT&RUN in CoREST1 KD cells that overexpressed CoREST2-FLAG. Gene ontology analysis of CoREST2-FLAG peaks revealed enrichment of peaks at genes associated with axon development and axonogenesis (Fig 7C and Extended Data Figure 10). Included among the genes with CoREST2 enrichment were *SOX4* and *ZNF800*, which both have reported roles in regulating EEC differentiation (Fig. 6D)^24, 25, 34^. Interestingly, there were several genes with enrichment of both STAT3 after ISX9 treatment and CoREST2, which included *SHH* and *INSM1* (Fig 6E). LSD1 was reported to demethylate the coiled-coil domain of STAT3, which is responsible for DNA binding^29, 30^. Knocking down LSD1 or CoREST2 attenuated the ISX9-induced increase in STAT3 chromatin binding (Fig 6F-6G). Altogether these results demonstrate that ISX9 increases the binding of STAT3 to genes important for EEC differentiation and that LSD1 and CoREST2 promote STAT3 chromatin binding.

**Figure 6.**
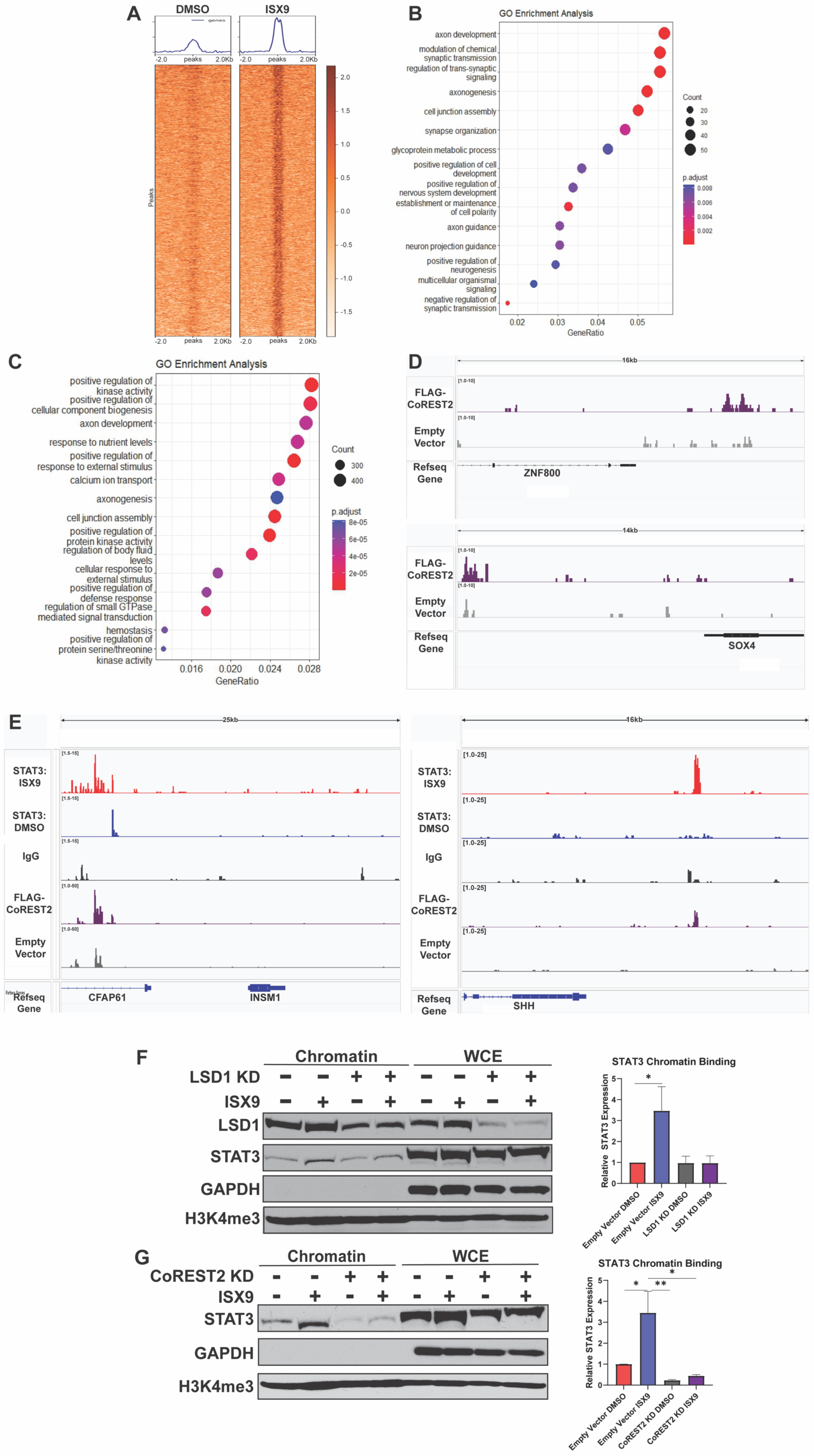
LSD1 and CoREST2 regulate STAT3 chromatin binding. **A.** Metagenomic heatmap for STAT3 CUT&RUN prepared from HT29 cells treated with DMSO or ISX9 (40 µM) for 4hrs. **B.** Gene Ontology enrichment analysis on genes associated with peaks from STAT3 CUT&RUN prepared from HT29 cells treated with ISX9. **C**. Gene Ontology analysis on peaks from FLAG CUT&RUN prepared from CoREST1 knockdown (KD) cells that overexpress CoREST2-FLAG. **D**. Gene tracks of CoREST2-FLAG enrichment upstream of *ZNF800* (top) and *SOX4* (bottom). Empty vector is a negative control in which cells contain two empty vector plasmids that do not express FLAG. **E**. Gene tracks showing enrichment of both STAT3 and CoREST2-FLAG upstream of *INSM1* (left) and *SHH (right).* **F.** Western blot of chromatin lysate prepared from empty vector cells and LSD1 KD cells treated with DMSO or ISX9 (40 µM) for 4hrs. Whole cell extract (WCE) serves as a control for chromatin extract. Graph shows results of densitometry quantification +/-SEM (N=5). **G.** Western blot of chromatin lysate prepared from empty vector cells and CoREST2 KD cells treated as in F. Graph shows results of densitometry quantification +/- SEM (N=3). Significance was determined one-way ANOVA with Tukey pairwise multiple comparison testing **(F and G).** *P≤ 0.05, ** P ≤ 0.01.

**Figure 7.**
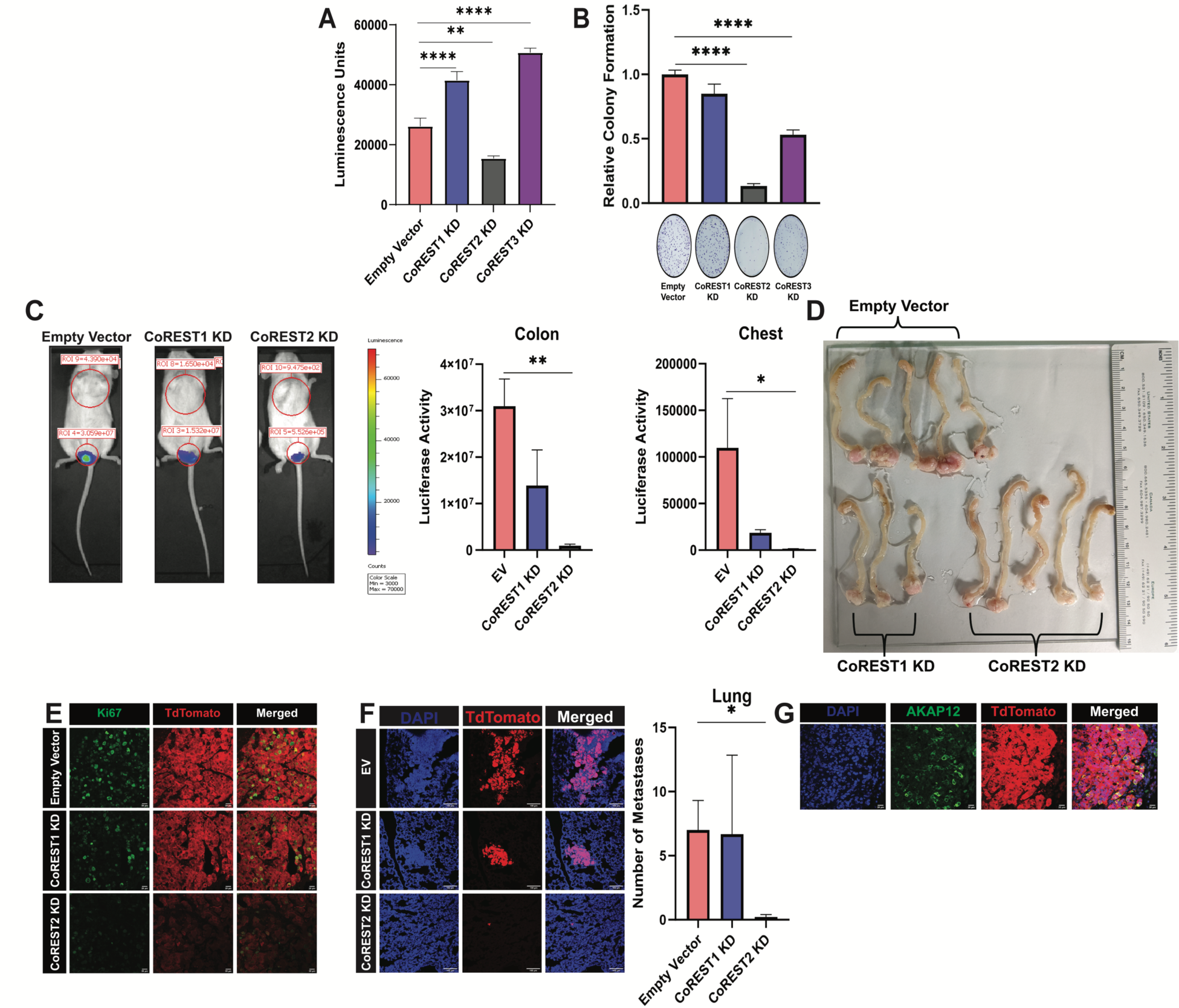
CoREST2 knockdown decr eases tumor growth and lung metastases. **A.** Cell viability of empty vector, CoREST1 knockdown (KD), CoREST2 KD, and CoREST3 KD HT29 cells. **B.** Clonogenic growth in empty vector, CoREST1 KD, CoREST2 KD, and CoREST3 KD HT29 cells. Graph is quantification the number of colonies per well relative to empty vector +/- SEM (N=6). **C.** Representative images of luciferase activity in colon and chest region of empty vector, CoREST1 KD, and CoREST2 KD HT29 cells orthotopically injected into colons of NSG mice at 2 weeks post-injection. Graphs show mean luciferase activity for each group in colon and chest regions +/- SEM (N=5 per group). **D.** Endpoint images of mouse colons with attached tumors at 3 weeks post-injection. **E.** Representative immunofluorescent images of Ki67 (green) and TdTomato^+^ (red, tumor cells) of colon tumor sections. **F.** Representative immunofluorescent images of TdTomato^+^ tumor cells (red) in lung tissue sections. Graph shows quantification of number of metastases formed +/- SEM (N=5 for empty vector and CoREST2 KD; N=3 for CoREST1 KD). **G.** Representative immunofluorescence images of ENLC marker AKAP12 (green) and TdTomato^+^ (red) of colon tumor from empty vector HT29 cells injected orthotopically into the colon of NSG mice. Significance was determined by one-way ANOVA with Dunnet pairwise multiple comparison testing **(A+B)** and Kruskal-Wallis with Dunn’s multiple comparison testing **(E)**. *P≤ 0.05, ** P ≤ 0.01, *** P ≤ 0.001, **** P ≤ 0.0001.

### CoREST2 Knockdown Decreases Tumor Growth and Lung Metastasis

EECs secrete factors to promote cancer cell survival^6^. Additionally, *NEUROG3* KD decreases cell viability (Extended Data Fig 8I-8J). These results indicate that targeting EEC differentiation may be an effective way to treat mCRC. To determine whether CoREST2 is a promising therapeutic target, cell viability was assessed after knockdown of individual CoREST paralogs. Interestingly, only CoREST2 KD decreased cell viability (Fig 7A). Additionally, CoREST2 KD dramatically decreased colony formation, while CoREST1 KD and CoREST3 KD had no and a moderate effect on colony formation, respectively (Fig 7B). To ascertain the effect of CoREST2 KD *in vivo,* we orthotopically implanted empty vector, CoREST1 KD, and CoREST2 KD HT29 cells that constitutively express luciferase and TdTomato into the distal colons of NSG mice. CoREST2 KD significantly reduced the primary colon tumor size, decreased the number of proliferative Ki67-positive cells in colon tumors, and reduced lung metastasis (Fig 7C-7F). In contrast, CoREST1 KD had a minimal effect on primary colon size, number of Ki67-positive cells, and metastasis (Fig 7C-7F). As tumor cell populations can differ *in vivo* versus *in vitro*, we performed immunofluorescence for AKAP12-positive ENLCs in the colon tumors. Surprisingly, ENLCs appeared to present at a higher percentage in the colon tumors than in vitro (Fig 7G). Collectively, this data suggests that CoREST2 is a promising target for treating mCRC.

## Discussion

Differences in cell state are a major source of tumor heterogeneity and contribute to aggressiveness and therapeutic resistance in cancer^1, 2^. Unlike non-mCRC, mCRC contains various differentiated cell types, including EECs, the neuroendocrine cell of the intestine^6^. EECs have previously been found to promote cancer cell survival in mCRC, akin to neuroendocrine cells promoting therapeutic resistance in other cancers^6, 10–13, 15, 35^. In this study, we sought to find factors that promote EEC differentiation in mCRC to identify new therapeutic targets for these difficult to treat cancers.

Cell differentiation in the colon is regulated by chromatin alterations and transcription factor activation^24, 25, 36, 37^. While many lineage-specifying transcription factors have been discovered, comparatively little is understood about the chromatin changes that accompany lineage determination in the normal colon or mCRC. It was revealed previously in the mouse duodenum, that there are changes in accessibility at subsets of enhancers in secretory progenitors compared to intestinal stem cells^36^. However, the study was unable to delineate chromatin changes between goblet cells and EECs, possibly due to reliance on bulk-ATAC sequencing of FACS-sorted cells. Our multi-omics data revealed that increased chromatin accessibility of EEC-promoting factors is important for EEC differentiation. Additionally, we identified the regulatory regions of EEC-promoting factors at which chromatin accessibility correlated the most with expression. Interestingly, we observed few regions of unique chromatin accessibility in other cell types. These findings suggest that EECs are unique in that dynamic chromatin changes occur during their differentiation.

Due to mutations causing altered signaling pathway activation in CRC, it is possible that CRCs contain additional cell populations not present in the normal colon epithelium. Through our multi-omics experiments, we identified a novel rare cell type with enteric neuron-like characteristics that we termed ENLCs. Evaluation of marker gene expression, RNA trajectory analysis, and diffusion mapping analysis showed ENLCs are distinct from the absorptive and secretory lineage. Despite their rarity *in vitro*, we observed an increase in these cells in our mouse model. Furthermore, our immunohistochemistry results demonstrated ENLCs are present in cancer patient samples but absent in the normal colon epithelium. These findings are interesting as the crosstalk between tumors and the peripheral nervous system through the secretion of neurotrophic factors and formation of synapses, has been linked to angiogenesis, immune cell regulation, and therapy resistance^38, 39^. Additionally, AKAP12, which was identified and used as an ENLC marker, has been associated with therapy resistance, increase in pro-tumor cancer-associated fibroblast, and metastasis in various cancers^40–42^. Our constructed GRNs indicate KLF6 and FOSL1 contribute to the differentiation of ENLC. Further studies are required to investigate the mechanism of ENLC differentiation and how ENLCs contribute to CRC.

Motif enrichment analysis of unique differentially accessible peaks in EECs identified many transcription factors with known and unknown functions in regulating EEC differentiation. STAT3 stood out as a potential EEC regulator, as STAT3 has previously been shown to promote the differentiation of other cell types through increasing expression of EEC transcription factor *NEUROG3*^26, 27^. STAT3 is also involved in neuroendocrine differentiation in prostate cancer and therapeutic resistance in CRC^28, 43^. Using knockdown and inhibitor-based strategies we demonstrated that STAT3 and upstream kinase JAK2 are regulators of EEC differentiation. In breast cancer, STAT3 has been shown to be activated by an increase in cytoplasmic calcium and ISX9-induced EEC enrichment is partly dependent on calcium signaling^20, 44^. Using a calcium chelator, we demonstrated that calcium activates STAT3 to promote EEC differentiation. Calcium-related signaling has previously been implicated in EEC differentiation in normal colon organoids and the Drosophila midgut, and neuroendocrine differentiation in breast and prostate cancer, suggesting that calcium signaling may generally drive neuroendocrine differentiation through STAT3 activation in normal and cancerous settings^20, 45–47^.

We previously published that LSD1 promotes EEC differentiation in mCRC, but without a clear mechanism^6^. It has previously been published that in response to interleukin-6, LSD1 can demethylate and promote STAT3 activity^30^. In this study, we demonstrate that LSD1 likely promotes EEC differentiation via demethylating STAT3. The enzymatic function of LSD1 is dependent on LSD1 being a part of a transcriptional regulatory complex, such as the CoREST complex. The core CoREST complex consists of LSD1, HDAC 1/2, and one of three CoREST paralogs^16–19^. Many studies have evaluated the biological roles of CoREST1, but few studies have investigated the function of CoREST2. Two prior studies have shown that CoREST2, encoded by the gene *RCOR2,* is upregulated in EEC progenitors in the normal colon^24, 48^, but whether there is a functional role for CoREST2 in EEC specification was not addressed. Our single cell RNA-seq, single cell multi-omics, and immunofluorescence data confirm that CoREST2 is primarily expressed in EECs. Additionally, we demonstrated that CoREST2 regulates STAT3 demethylation in an LSD1-dependent manner. The coiled-coil domain of STAT3 mediates STAT3 DNA binding and was reported as the target of LSD1 mediated-demethylation^29, 30^. In line with this, we revealed that LSD1 and CoREST2 impact chromatin binding of STAT3. Also of note, our constructed GRN showed that NEUROG3 regulates CoREST2 expression in EECs suggesting the possibility that the increased expression of CoREST2 acts in a positive feedback loop to maintain the expression of EEC genes such as *NEUROG3* and *INSM1* through interacting with STAT3. In summation, our data supports a model in which LSD1 and CoREST2 interact with STAT3 to enhance or prolong the binding of STAT3 to genes important for EEC differentiation, likely through STAT3 demethylation.

Our CUT&RUN data showed that both STAT3 and CoREST2 localize to genes associated with neuronal differentiation and functions. EECs share many of the same transcription factors as neurons and morphologically some EECs have synapse-like neuropods^49^, making many of these GO pathways related to EECs. Additionally, we identified genes with enrichment of both STAT3 and CoREST2. As many differentiation pathways and molecules are shared between EECs, neuroendocrine cells, and neurons our findings related to the interaction of LSD1-CoREST2 and STAT3 will likely be applicable to other cell and cancer types.

Finally, we demonstrated that targeting CoREST2, in an orthotopic xenograft mouse model, significantly decreases primary tumor formation in the colon and metastasis to the lung. There are several clinically relevant LSD1 inhibitors, which were all optimized to inhibit LSD1 demethylation of lysine 4 of histone H3. These inhibitors have significant associated toxicities because LSD1 is critical for normal biological functions such as hematopoiesis^50, 51^. Our findings regarding LSD1 promoting EEC differentiation in conjunction with STAT3 and CoREST2 suggest that designing LSD1 inhibitors targeting LSD1-CoREST2 and/or the interaction of LSD1 with STAT3 may have improved efficacy in mCRC.

## Methods

### Cell lines and organoids growth and treatment

All cell lines were maintained in a 37℃ humidified incubator with 5% CO_2_. HT29 cells were cultured in McCoy 5A media (Corning), NCI-H508 and SW403 cells in RPMI1640 media (Corning), and LS174T cells in DMEM, supplemented with 10% FBS (Gibco). All cell lines were purchased from ATCC and authenticated and tested for Mycoplasma using the Universal mycoplasma detection kit (ATCC, 30-1012K). Normal human colon organoids derived from the ascending colon (83) were obtained from Dr. Jason Spence and the University of Michigan Translational Tissue Modeling Laboratory. Organoids were passed in Intesticult organoid growth media (STEMCELL # 06010) and differentiated in Intesticult organoid differentiation media (STEMCELL # 100-0214). ISX9 (Tocris #4439), Stattic (Selleckchem #S7024), S31-201 (Selleckchem #S1155), Corin (MedChemExpress #HY-111048), BAPTA-AM (APExBio #B4758), AZD1460 (Selleckchem #S2162), and Ruxolitinib (Selleckchem # S1378) were solubilized in DMSO (VWR #97063-136) prior to treatment.

### Single Cell Multi-omics

Approximately 10,000 cells per sample were sequenced via the 10X Genomics Chromium System using the Chromium Next GEM Single Cell Multiome ATAC + Gene Expression Kit A at the Indiana University School of Medicine (IUSM) Center for Medical Genomics core. The libraries were sequenced at the IUSM Center for Medical Genomics using a NovaSeq 600 with approximately 375 million ATAC read pairs and 500 million Gene Expression read pairs per sample.

### Single-cell data pre-processing and QC

Read alignment and gene-expression quantification of HT29 cells scRNA-seq and scATAC-seq data was performed using the CellRanger-arc Count pipeline (version 2.0.2, 10X Genomics). The CellRanger pre-built human reference package was used for read alignment (hg38). The filtered feature matrices output was then used to create a Seurat object using the Seurat package v4.3.0.1^52^. For scRNA-seq, cells were filtered to include only cells with no more than 25% mitochondrial gene. The data were normalized, and highly variable genes were identified and scaled using *SCTransform*. Next, dimensionality reduction by principal components (PCs) was calculated using *RunPCA* and to estimate the significant number of PCs to be used *ElbowPlot* function was used. Next, the uniform manifold approximation and projection (UMAP) embedding were calculated and visualized using *RunUMAP* and *DimPlot*. Unsupervised Louvain clustering was carried out using *FindNeighbors* and *FindClusters*. In our analysis, we used Seurat v4.3.0.1 to perform batch-effect correction. 3000 highly variable genes were defined within the 2 samples (ISX9 and control) with the Seurat *FindVariableFeatures* function. We also identified unsupervised integration “anchors” for similar cell states using shared nearest neighbor graphs (*FindIntegrationAnchors*), and then integrated our 2 different datasets using these anchors using *IntegrateData*.

For scATAC-seq, data was processed using the standard Signac workflow^53^. Cells with fewer than 300 or more than 100,000 ATAC fragments were filtered, as were cells with nucleosomal enrichment > 1.5 or transcriptional start site enrichment < 1. Peaks within each dataset were identified using MACS2. Genomic blacklist regions were removed using *subsetByOverlaps.* Dimensional reduction was performed with latent semantic indexing (LSI) via the *RunTFIDF* command, *FindTopFeatures* function, and *RunSVD* function. UMAP projection was performed utilizing LSI components 2-50. The two scATAC-seq datasets were integrated using Harmony package^54^. RNA-seq data were integrated with ATAC-seq data by constructing weighted nearest neighbor (WNN) graph using *FindMultiModalNeighbors*.

### Cluster annotations and data visualization

To identify different major cell populations, we manually assigned class identities based on the expression of well-established marker genes. Some clusters were excluded from the analysis because they expressed either apoptotic markers or interferon-related genes. Gene markers for each cluster were identified using the Seurat *FindAllMakers* command with the following settings: min.pct = 0.25, test.use = ‘‘wilcox’’, only.pos = TRUE, logfc.threshold = 0.25. Clusters were manually annotated based on the expression of well-established marker genes.Transcription factor motif enrichment was implemented with the ChromVAR software package implemented through Signac. JASPAR 2020 vertebrate transcription factor motifs were utilized. ChromVAR results were imported to Signac as an assay object. ATAC peaks linked with gene expression were calculated by using the Signac *LinkPeaks* command with default settings.

### RNA velocity

Spliced/unspliced expression matrices were generated as loom files using Velocyto^55^. Seurat objects were converted into AnnData objects containing the corrected counts, clusters, and UMAP embeddings. Then, the loom files were merged with the AnnData objects and loaded into scVelo (v0.2.1)^56^, the ratio of spliced to unspliced reads per cluster was found, and cell velocities were computed. All functions were run with default settings unless otherwise stated. The *scvelo.pp.filter_and_normalize* argument ‘n_top_genes’ was set to 3000, and the ‘n_npcs’ and ‘n_neighbors’ arguments of *scvelo.pp.momentum* were both set to 30. The velocity cell arrows were made with the *scvelo.pl.velocity_embedding* function. The top velocity genes per cluster were discovered using *scvelo.tl.rank_velocity_genes*, and plotted using *scvelo.pl.velocity*. *scvelo.tl.velocity_confidence* generated the velocity confidence and length values, and the results were plotted using *scvelo.pl.scatter*. The RNA-velocity analysis was extended through calculating RNA splicing kinetics using dynamic model using *scv.tl.recover_dynamics* and scv.tl.*velocity*(mode=’dynamical’). Cluster-specific identification of potential drivers was discovered using *scv.tl.rank_dynamical_genes*.

### Simulated gene perturbation of ISX9 sample

Simulated gene perturbation was performed using CellOracle^57^. We constructed our own gene-regulatory networks (GRN) from our scATAC-seq data using Cicero package. First, our scATC-seq data were converted to cicero object and cis-regulatory interactions were identified using *run_cicero.* Transcription start sites (TSS) were annotated using hg38 genome as reference using CellOracle. TSS information was integrated with cis-regulatory interactions using *integrate_tss_peak_with_cicer.* Peaks with weak coaccessibility scores using *integrated*.*coaccess* >= 0.8 command. Transcription factor motif scan was performed using *tfi.scan*. the final GRN and 3000 highly variable gene expression from ISX9 gene matrix were loaded into the CellOracle object using *co.import_TF_data* and *co.import_anndata_as_raw_count*, respectively. We constructed a cluster-specific GRN for all clusters using *oc.get_links* and kept only network edges with p-value <=0.01. To simulate gene overexpression or knockout, we perturbed the gene expression to 1.5 or 0, respectively in the *oc.simulate_shift* function.

### Generation of stable cell lines

For empty vector (VectorBuilder), CoREST2-FLAG overexpression (VectorBuilder) and knockdown of STAT3 (Sigma, SHCLNG-NM_003150, TRCN0000329887), JAK2 (Sigma, SHCLNG-NM_004972, TRCN000002181), CoREST1 (Sigma, SHCLNG-NM_015156, TRCN0000418894), CoREST2 (Sigma, SHCLNG-NM_17585, TRCN0000122645), CoREST3 (Sigma, SHCLNG-NM_018254, TRCN0000116089), LSD1 (Sigma, SHCLNG-NM_015013, TRCN0000046071), NEUROG3 (Sigma, SHCLNG-NM_020999, TRCN0000427521) and empty vector TRC2 (Sigma, #SHC201), the following lentiviral protocol was used. Briefly, 2.5×10^5 HEK293T cells were plated on day 1 in DMEM 1X containing 10% FBS. The next day cells were transfected with shRNA of interest, EV control, and packaging plasmids. On the third day, the media was replaced with fresh DMEM with 10% FBS. 24hrs later, media containing lentiviral particles was collected and filtered using a 0.22 µm filter, and concentrated with Lenti-X^TM^ Concentrator (Takara, #631232). Media was replaced and on day 5, the media was again collected, filtered using a 0.22 µm filter, and concentrated with Lenti-X^TM^ Concentrator and combined with the media collected on day 4. To perform the transduction, virus-containing media and polybrene were added to cells. The next day, to select for transduced cells, cells were treated with hygromycin (400 µg/mL) (Millipore # 400052) or puromycin (2µg/mL) (Sigma-Aldrich #P8833).

### Immunofluorescence and imaging

In cell lines, for AKAP12, βIII-Tubulin, CoREST2, and INSM1 immunofluorescence (IF) was performed as previously described^58^. Cells were incubated with anti-AKAP12 (Thermo Fisher #PA5-21759, 1:200), anti-βIII-Tubulin (R&D Systems, MAB1195-SP, 1:500), anti-CoREST2 (Sigma-Aldrich, #HPA021638, 1:200), or anti-INSM1 (Santa Cruz Biotechnology #271408, 1:100) in 1% BSA in PBST for 1 hour at room temperature. For STAT3 IF, cells were stained with anti-STAT3 (CST #9139, 1:800)) according to manufacturer’s protocol. Cells were fixed with 4% paraformaldehyde, permeabilized with methanol, and incubated overnight at 4°C with anti-STAT3 antibody in 1% BSA/PBST. For AKAP12, βIII-Tubulin, CoREST2, and STAT3 cells/tissue were then incubated in goat anti-mouse IgG Alexa Fluor 488 (CST #4408, 1:1000), goat anti-mouse IgG Alexa Fluor 594 (CST # 8890, 1:500) or goat anti-rabbit IgG Alexa Fluor 594 (CST #8889, 1:500) in 1% BSA in PBST for 1 hour at room temperature. Images were acquired on a Leica SP8 scanning confocal system with MDi8-inverted microscope with LASX software (Leica Microsystems). All the images were taken at either X40 or X63 magnification, 1.4A oil immersion at room temperature, and processed using ImageJ.

### Immunohistochemistry (IHC)

AKAP12 was detected by IHC on a tissue colon carcinoma tissue microarray (TissueArray #CO20001A) following unmasking in Tris-EDTA buffer. Anti-AKAP12 antibody was applied at a 1:500 dilution followed by anti-rabbit HRP (CST # 8114) and DAB substrate (CST # 8059). Cells were counterstained in hematoxylin. Cores were scored based on intensity of staining and percentage of cells with positive staining: Samples with no or little tumor sample were excluded.

### RNA isolation and RT-qPCR

From cell lines, total RNA was isolated from cell pellets using the RNeasy mini kit (Qiagen #74104) according to the manufacturer’s protocol. For organoids, total nucleic acids were first isolated from cell pellets first using Trizol Reagent (Ambion #15596026) and chloroform. The nucleic acid fraction was then added to the column of a RNeasy microkit (Qiagen # 74004) and the manufacturer’s protocol was followed. The Maxima first strand cDNA synthesis kit (Thermo Fisher #K1642) was used to synthesize cDNA. For quantitative reverse transcription PCR, cDNA was amplified using gene-specific primers and FastStart Essential DNA Green Master (Roche #06402712001). Cq values of genes of interest were normalized to housekeeping gene *RHOA* expression.

### Whole-cell isolation and western blot analysis

For whole cell protein isolations, cell pellets were lysed in 4% SDS using a qiashredder. Analysis of relative densitometry of Western blots was determined using ImageJ software and normalized to density of H3K4me3 loading control. Results are shown as mean ± SEM.

### Nuclear Immunoprecipitations (IPs)

Nuclear IPs were performed as previously described^59^. Briefly, 3.5×10^6, 4.0×10^6, or 7.0×10^6 cells were cultured in 150 mm plates for approximately 48hrs for HT29, NCI-H508, and SW403 cells, respectively. Cell pellets were used to perform nuclear extraction using CEBN Buffer [10 mmol/L HEPES, pH 7.8, 10mmol/L KCl, 1.5 mmol/L MgCL_2_, 0.34 mol/L sucrose, 10% glycerol, 0.2% NP-40, 1X protease inhibitor cocktail (Sigma, #P5726) and 1X phosphatase inhibitor (Thermo, #88266)] and then washed with CEB buffer (CEBN buffer minus NP-40) containing both inhibitors. To extract the nuclear fraction, cells pellets were then resuspended in modified RIPA buffer (50nm Tris pH 7.5, 150nM NaCl, 5mM EDTA, 50nM NaF, both inhibitors) and sonicated using Bioruptor® Pico (Diagenode). To remove the possibility of any DNA-dependent protein interactions, the nuclear fractions were incubated with spermine (50nM) and spermidine (15mM) for 1 hour on a rotator at 4°C. The nuclear extract was then rotated with antibody and protein A/G coated magnetic beads overnight at 4°C. The following day, beads were washed in TNE buffer, (50nM Tris pH 7.5, 150nM NaCl, 5mM EDTA, 50nM NaF, 0.5% NP-40 and 0.5% TritonX-100) protein was eluted and analyzed by western blot.

### CUT&RUN

Cleavage Under Targets & Release Using Nuclease (CUT&RUN) was performed according to the CUTANA CUT&RUN protocol. In summary, 5×10^6 live cells were washed and resuspended in wash buffer (20 mM HEPES, pH 7.5, 150 mM NaCl, 0.5mM Spermidine and 1X Protease Inhibitor). Next, cells were tethered to activated ConA magnetic beads (CUTANA^TM^ Concanavalin A, #21-1401) which were washed and resuspended in bead activation buffer (20mM HEPES, ph 7.9, 10mM KCl, 1mM CaCl_2_. 1mM MnCl_2_). Cells were then permeabilized in antibody buffer (wash buffer + 0.01% digitonin + 2mM EDTA) and incubated with STAT3 (CST, #9139) or FLAG antibody (Sigma Aldrich #F3165) on nutator overnight at 4°C. Cell-bead slurry was then washed twice with cold digitonin buffer (wash buffer + 0.01% digitonin), incubated with pAG-MNase (Epicypher, #15-1116) for 10 min at RT, and then washed twice more with cold digitonin buffer. MNase was then activated by the addition of CaCl_2_ to cleavage targeted chromatin for 2 hours at 4°C. Following chromatin digestion, MNase activity was stopped, and chromatin fragments were released into supernatant by adding stop buffer (340 mM NaCl, 20mM EDTA, 4mM EGTA, 50 µg/mL RNase A, and 50 µg/mL Glycogen) and incubating at 37°C for 10 min. DNA was purified from the collected supernatant using CUTANA^TM^ DNA Purification Kit (EpiCypher #14-0050) per manufacturer’s instruction. Finally, 5 ng of purified CUT&RUN enriched DNA was used to prepare Illumina library using the NEBNext Ultra II DNA Library Prep kit (NEB, #E76450) per the manufacturer’s protocol, followed by sequencing.

### CUT&RUN Sequencing analysis

Sequencing read quality control for all samples was assessed with FastQC (v0.12.1). CUT&RUN sequences were aligned to the hg38 reference genome using Bowtie2 (v2.5.1). Peaks were called with MACS (v2.2.7.1). Heatmaps and Counts Per Million normalized bigWigs were created using deepTools (v3.5.1) bamCoverage, computeMatrix,and plotHeatmap. IGV was utilized for creating gene tracks. Gene ontology analysis was performed using ChIPseeker (v1.38.0) and ClusterProfiler (v4.10.0).

### Chromatin Extraction

3.5×10^6 cells were cultured in 150 mm plates for approximately 48hrs. First cell pellets were used to perform nuclear extraction using CEBN buffer and was then washed with CEB buffer (both containing inhibitors). Soluble nuclear extract was then obtained by incubating nuclear extract in soluble nuclear buffer (2mM EDTA and 2mM EGTA, plus inhibitors) for 30 min on a rotator at 4°C. The remaining cell pellet is the total chromatin fraction, which was then lysed using a qiashredder (Qiagen) and 4% SDS and analyzed by Western blot.

### Clonogenic growth assay

For clonogenic growth assays, 500 single cells were plated in one well of a 6-well plated and allowed to grow for 14 days at 37°C. Cells were then fixed with ice-cold methanol for 10 minutes and stained with crystal violet. Results were analyzed using ImageJ. Results are shown as mean +/- SEM.

### Cell Titer-Glo Assay

Viability assays were performed using the CellTiter-Glo Luminescent Cell Viability Assay (Promega #G7572) per manufacturer’s protocol. Briefly, 1.0 × 10^3 cells were plated in 96-well plates and allowed to incubate under standard growth conditions. Results are shown as mean +/- SEM.

### Orthotopic xenograft mouse model

HT29 cells previously infected with pLenti-U6-tdTomatato-P2A-BlasR (LRT2B) virus (Addgene, 1108545) that enables dual production of firefly luciferase and TdTomato were used. 2.0×10^5 empty vector, CoREST1 KD, or CoREST2 KD cells were combined with Matrigel and injected into the mechanically prolapsed mouse large intestine. After approximately 3 weeks, mice were sacrificed and xenograft primary tumors, along with adjacent normal mouse tissue, lungs and livers were harvested, fixed in 4% paraformaldehyde, and frozen in optimal cutting temperature media for further analysis. *In vivo* imaging was done using an IVIS imaging system. For IF, tissue was thawed, blocked for 1 hour in 1% BSA in PBST with 5% normal goat serum, and incubated overnight at 4°C in a humified chamber with anti-AKAP12 antibody (Thermo Fisher #PA5-21759, 1:200) or anti-Ki67 (CST #9449, 1:1000) in 1% BSA in PBST. The next day tissue was incubated with goat anti-rabbit IgG Alexa Fluor 488 (CST #4412, 1:1000) in 1% BSA at room temperature. All mouse experiments were covered under a protocol approved by the Indiana University Bloomington Animal Care and Use Committee in accordance with the Association for Assessment and Accreditation of Laboratory Animal Care International.

## Antibodies

LSD1 (CST: 2139) (Western blot), LSD1 (CST:4218) (IP), STAT3 (CST:9139) (Western blot, IF, CUT&RUN), CoREST2 (Sigma: HPA021638) (IF), Neuron-specific βIII-Tubulin (R&D Systems: MAB1195-SP) (IF), CoREST (CST: 14567)(Western blot) JAK2 (CST: 3230) (Western blot) AKAP12 (Thermo Fisher: PAS-21759) (IF), INSM1 (SCB: 271408) (IF) FLAG (Sigma: F1804) (Western blot, IP, CUT&RUN), P-STAT3^Y705^ (CST:9145) (Western blot), Histone H3 (CST: 9717) (Western blot), Tri-Methyl Histone H3 (Lys4) (CST: 9751) (Western), Di-Methyl Lysine Motif (CST: 14117) (IP), β-actin (CST: 8457) (Western blot), GAPDH (CST: 5714) (Western), Mouse IgG2a (CST:61656)(IP, CUT&RUN), Rabbit IgG (Millipore 12370) (IP)

### qPCR primers

*RHOA* forward: CGTTAGTCCACGGTCTGGTC

*RHOA* reverse: ACCAGTTTCTTCCGGATGGC

*INSM1* forward: ACATCAACAAGTGCCACCCA

*INSM1* reverse: CGCACTCTCTTTGTGGGTCT

*NEUROG3* forward: CTCACCAAGATCGAGACGCT

*NEUROG3* reverse: GTACAAGCTGTGGTCCGCTA

*ATOH1* forward: AGAGAGCATCCCGTCTACCC

*ATOH1* reverse: GCTCCGGGGAATGTAGCAAA

*LGR5* forward: AGACACGTACCCACAGAAGC

*LGR5* reverse: AACGCATTGTCATCCAGCCA

*HES1* forward: AAAAATTCCTCGTCCCCGGT

*HES1* reverse: GGCTTTGATGACTTTCTGTGCT

*MUC2* forward: GCTATGTCGAGGACACCCAC

*MUC2* reverse: AGACGACTTGGGAGGAGTTG

*SPDEF* forward: CTCAGCTGCCCACACCTCTT

*SPDEF* reverse: GGGATACGCTGCTCAGACC

*GFI1* forward: CTCGCCCACCTCTTCCAAATTTAAC

*GFI1* reverse: GTCACTCCGAGGGCTTGCTC

*RCOR1* forward: AAGACGCAGTCAAGAACGGG

*RCOR1* reverse: TGCCAGAAGAGCATCCCAAG

*RCOR2* forward: TCAGCTCATCTCCCTCAAGC

*RCOR2* reverse: CAGCGGGAGTTGAACTTGGT

*RCOR3* forward: CCGAGTTACTGGGGAAGAACC

*RCOR3* reverse: TCCCAACATCGTGCTCGTC

## Acknowledgements

We would like to thank the Indiana University School of Medicine (IUSM) Center for Medical Genomics core for their assistance with performing the scRNA-seq, the IU Center for Genomics and Bioinformatics for their assistance with library preparation and sequencing of CUT&RUN samples, and the IU Bloomington Light Microscopy Imaging Center for maintaining and providing access to imaging equipment. This work was in part supported by a Research Enhancement Grant [to H. M. O’Hagan] from the Indiana University School of Medicine (IUSM), a Core Pilot Grant [to H. M. O’Hagan] from the Indiana Clinical and Translational Sciences Institute funded, in part by Grant Number UL1TR002529 from the National Institutes of Health, National Center for Advancing Translational Sciences, Clinical and Translational Sciences Award. and pilot funding [to H. M. O’Hagan] from the IU Simon Comprehensive Cancer Center (IUSCCC) Tumor Microenvironment & Metastasis Program and the IUSCCC P30 Support Grant (P30CA082709). Preliminary data was also funded in part by the IUSCCC Joe Ward Fellowship [to C. Ladaika]. The content is solely the responsibility of the authors and does not necessarily represent the official views of the NIH or IUSM. Additional pilot funding was provided by the NCI SPORE Project, Epigenetic Therapies – New Approaches (P50CA254897) Developmental Research Program [to H. M. O’Hagan]. A. Ghobashi and C. Ladaika were supported by the Doane and Eunice Dahl Wright Fellowship generously provided by Ms. Imogen Dahl.

## Extended Data Inventory

**Extended Data Figure 1. Single cell multi-omics cluster gene expression (excel file).**

**Extended Data Figure 2. Additional feature linkage maps**.

**Extended Data Figure 2.**
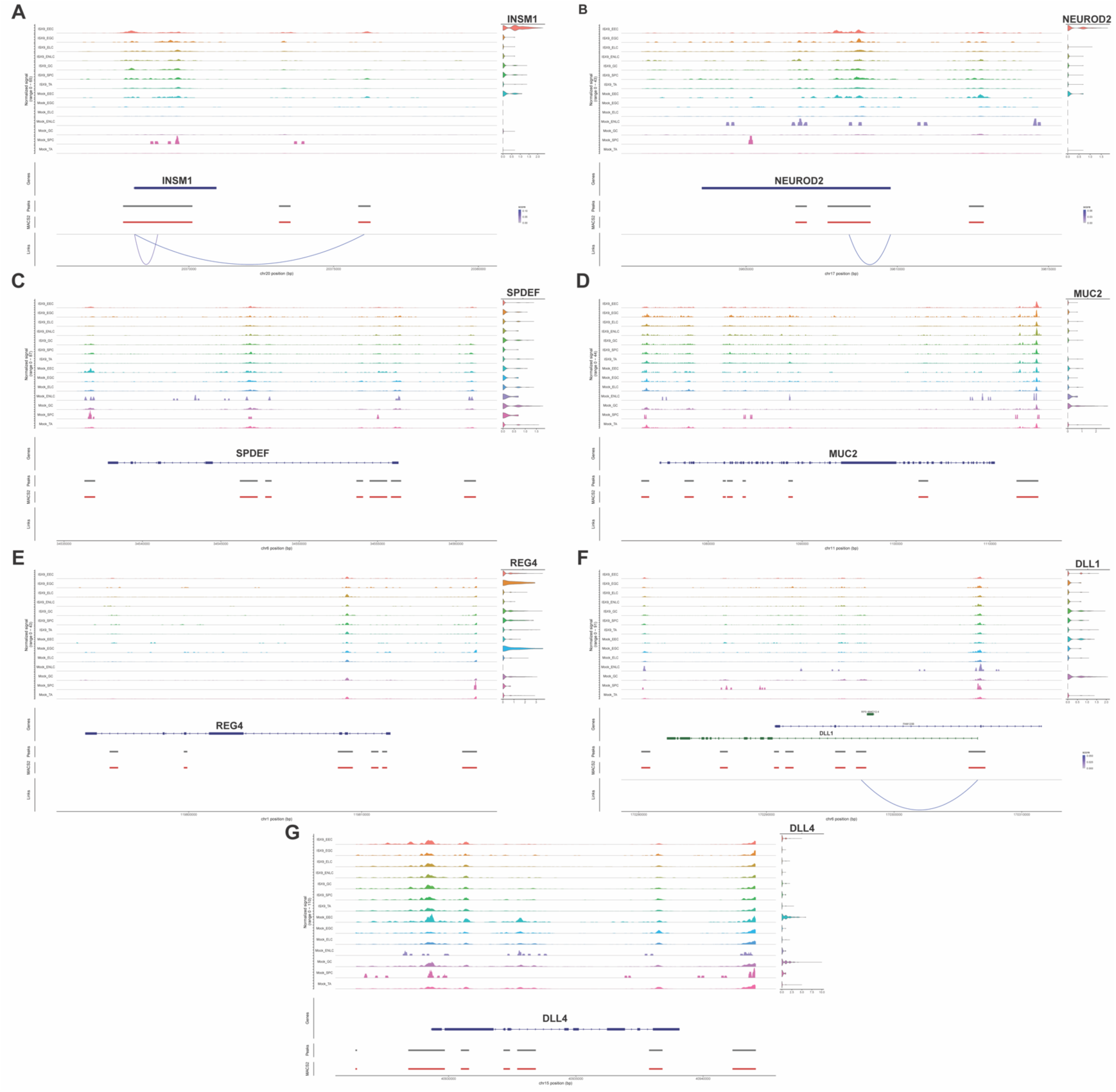
Additional feature linkage maps. **A-B.** Peaks to genes linkage maps of EEC transcription factors *INSM1* and *NEUROD2*. Peaks show ATAC-seq peak accessibility at *INSM1* and *NEUROD2* regulatory regions, violin plots show expression of *INSM1* and *NEUROD2,* curved blue lines show significant correlation between peak accessibility and expression of *INSM1* and *NEUROD2*. **C-D.** Peaks to genes linkage maps of goblet cell markers *SPDEF* and *MUC2*. Peaks show ATAC-seq peak accessibility at *SPDEF* and *MUC2* regulatory regions, violin plots show expression of *SPDEF* and *MUC2. **E.*** Peaks to genes linkage map of early goblet cell marker *REG4*. Peaks show ATAC-seq peak accessibility at *REG4* regulatory regions and violin plot shows expression of *REG4*. **F-G.** Peaks to genes linkage maps of EEC transcription factors *DLL1* and *DLL4*. Peaks show ATAC-seq peak accessibility at *DLL1* and *DLL4* regulatory regions, violin plots show expression of *DLL1* and *DLL4,* curved blue lines show significant correlation between peak accessibility and expression of *DLL1* and *DLL4*.

**Extended Data Figure 3. Single cell multi-omics differentially accessible peaks (excel file).**

**Extended Data Figure 4. Gene regulatory network data (excel file).**

**Extended Data Figure 5.**
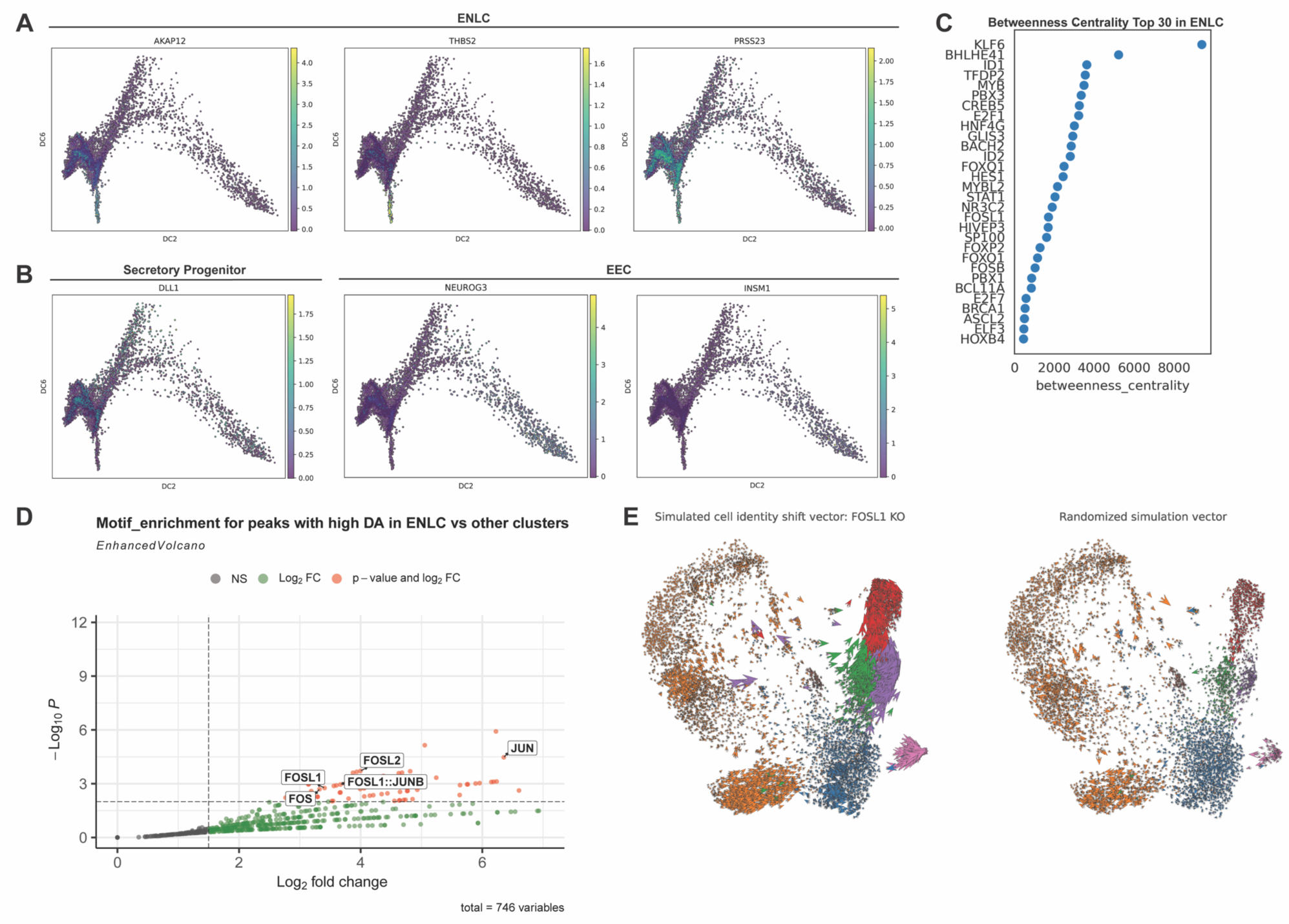
*In silico* simulation of FOSL1 KO on EEC and ENLC specification. **A.** Expression of ENLC marker genes *AKAP12*, *THBS2*, and *PRSS23* in cells arranged in diffusion pseudotime. **B.** Expression of secretory progenitor marker gene *DLL1* and EEC marker genes *NEUROG3* and *INSM1* arranged in diffusion pseudotime. **C.** Top 30 transcription factors based on betweenness centrality from ENLC gene regulatory network (GRN). **D.** Volcano plot showing transcription factor motif enrichment in differential accessible peaks in EECs vs non-EEC clusters. Red dots have a log2FC > 1.5 and p-value < 0.05. **E.** RNA velocity plot showing cell identity shift following *FOSL1* simulated knockout (left) and randomized simulated vector (right).

**Extended Data Figure 6.**
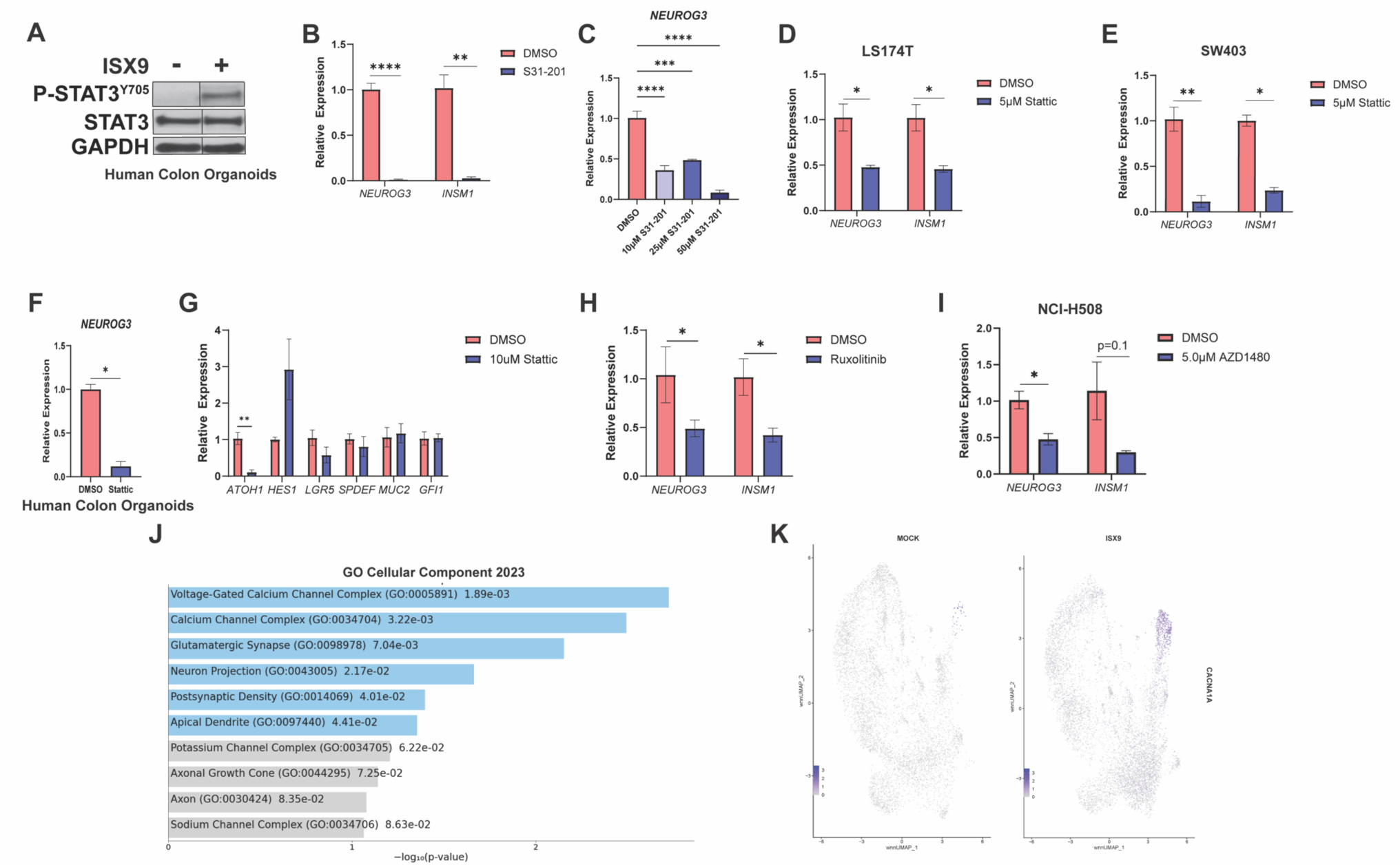
STAT3, JAK2, and calcium promote EEC differentiation. **A.** Western blot of human colon organoids treated with vehicle or ISX9 (40 µM) for 48hrs, plus one additional 24hr treatment. Black line indicates lanes were removed between displayed lanes. **B.** qRT-PCR of EEC markers after treating HT29 cells twice for 24hrs with vehicle or STAT3 inhibitor (S31-201, 100 µM). **C.** qRT-PCR of EEC markers after treating HT29 cells twice for 24hrs with 10 µM, 25 µM, or 50 µM S31-201. **D.** qRT-PCR of EEC markers after treating LS174T cells twice for 24hrs with vehicle or Stattic (5 µM). **E.** qRT-PCR of EEC markers after treating SW403 cells twice for 24hrs with vehicle or Stattic (5 µM). **F.** qRT-PCR of *NEUROG3* after treating human colon organoids cells twice for 48hrs with vehicle or Stattic (10 µM). **G.** qRT-PCR of secretory cell marker *ATOH1*, absorptive cell marker *HES1*, stem cell marker *LGR5*, and goblet cell markers *SPDEF, MUC2,* and *GFI1* after treating HT29 cells twice for 24hrs with vehicle or Stattic (10 µM). **H.** qRT-PCR of EEC markers after treating HT29 cells twice for 24hrs with vehicle or JAK2 inhibitor Ruxolitinib (5 µM). **I.** qRT-PCR of EEC markers after treating NCI-H508 cells twice for 24hrs with vehicle or JAK2 inhibitor AZD1408 (5 µM). **J.** Gene Ontology analysis of top 100 genes differential expressed in EEC cluster relative to all other clusters based on adjusted p-value. **K.** Feature plot of normalized expression values of CACNA*1A.* Significance was determined by Student’s t-test **(B and D-I)** and one-way ANOVA with Dunnet pairwise multiple comparison testing **(C).** *P≤ 0.05, ** P ≤ 0.01, *** P ≤ 0.001, **** P ≤ 0.0001.

**Extended Data Figure 7.**
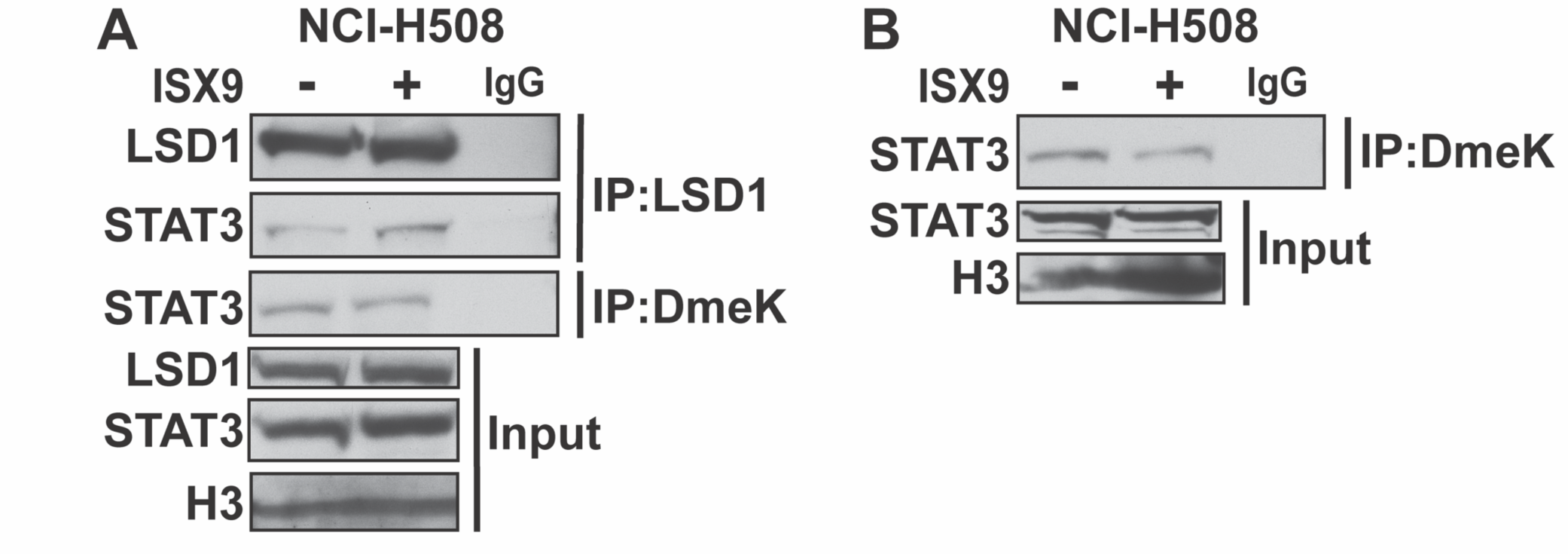
LSD1 Interacts with STAT3 and STAT3 is demethylated in NCI-H508 cells. **A.** Western blot of LSD1 immunoprecipitation (IP) and Dimethyl-lysine (DmeK) IP performed using nuclear lysates prepared from NCI-H508 cells treated with vehicle or ISX9 (40 µM) for 4hrs. IgG IP serves as a negative control. Western blot of DmeK IP performed using nuclear lysates prepared from NCI-H508 cells treated with vehicle or ISX9 (40 µM) for 4hrs. IgG IP serves as a negative control.

**Extended Data Figure 8.**
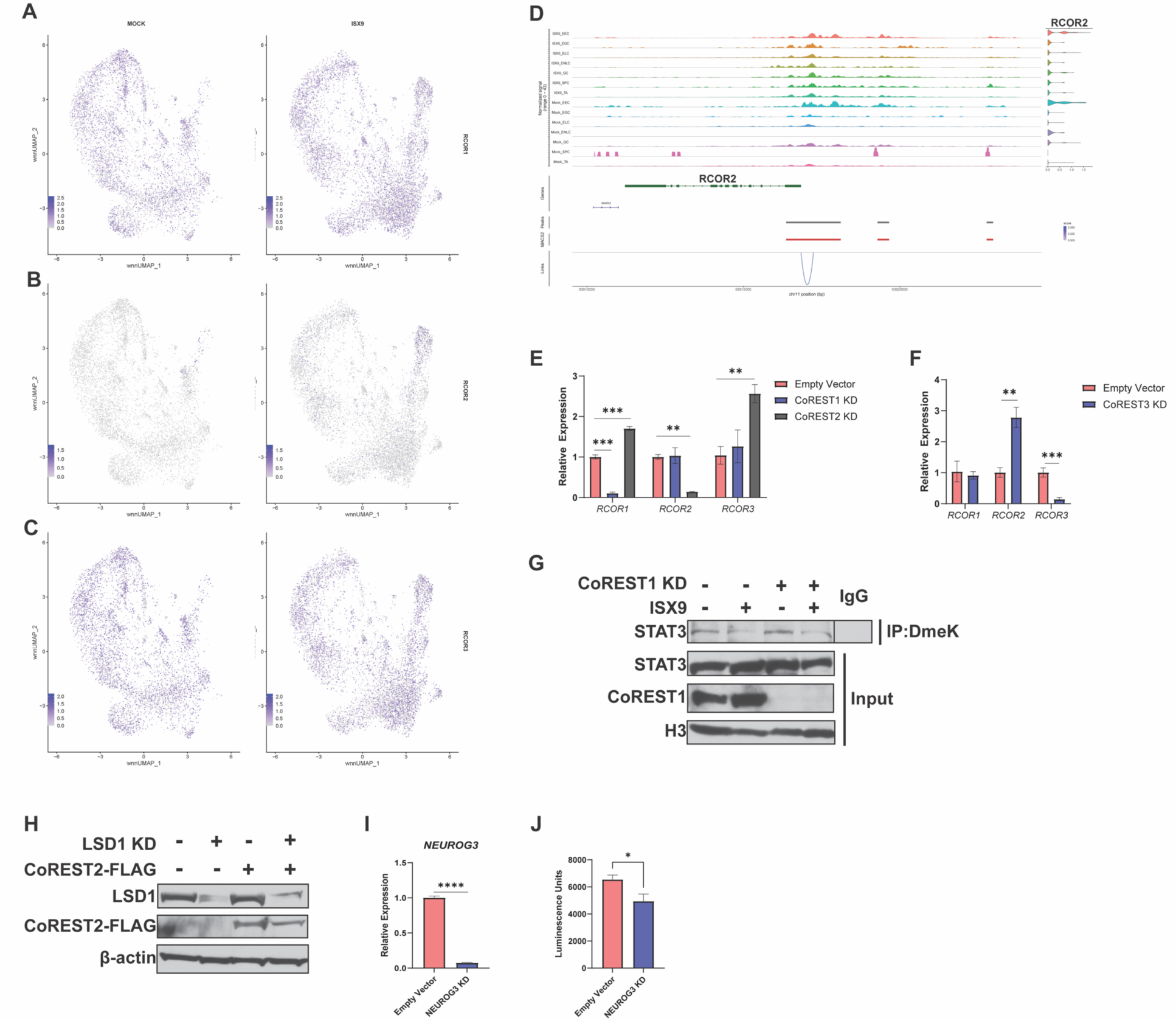
CoREST2 is stabilized by LSD1. **A-C.** Feature blots of normalized expression values of *RCOR1, RCOR2,* and *RCOR3.* **D.** Peaks to genes linkage map of *RCOR2*. Peaks show ATAC-seq peak accessibility at *RCOR2* regulatory regions, violin plot shows expression of *RCOR2*, curved blue line shows significant correlation between peak accessibility and expression of *RCOR2*. **E.** qRT-PCR in empty vector, CoREST1 KD, and CoREST2 KD HT29 cells**. F.** qRT-PCR in empty vector and CoREST3 KD HT29 cells. **G.** Western blot of Dimethyl-lysine (DmeK) IP performed using nuclear lysates prepared from empty vector and CoREST1 KD HT29 cells treated with vehicle or ISX9 (40 µM) for 4hrs. IgG IP serves as a negative control. Black line indicates lanes were removed between displayed lanes. **H.** Western blot of empty vector and LSD1 KD transduced with an empty vector or CoREST2 OE FLAG plasmid. **I.** qRT-PCR in empty vector and NEUROG3 KD HT29 cells. **J.** Cell viability of empty vector and NEUROG3 KD cells. Significance was determined by one-way ANOVA with Dunnet pairwise multiple comparison testing **(E)** and Student’s t-test **(F, I, and J)**. *P≤ 0.05, ** P ≤ 0.01, *** P ≤ 0.001, **** P ≤ 0.0001.

**Extended Data Figure 9. STAT3 CUT&RUN (excel file)**. Called peaks from STAT3 CUT&RUN from HT29 cells treated with DMSO or ISX9 (40 µM) for 4hrs. IgG CUT&RUN served as a negative control.

**Extended Data Figure 10. CoREST2-FLAG CUT&RUN (excel file).** Called peaks from FLAG CUT&RUN prepared from CoREST1 knockdown cells that overexpress CoREST2-FLAG. Cells that contain two empty vector plasmids that do not express FLAG were used as a negative control.

## Notes

### Competing Interest Statement

The authors have declared no competing interest.

